# Distributed control circuits across a brain-and-cord connectome

**DOI:** 10.1101/2025.07.31.667571

**Authors:** Alexander Shakeel Bates, Jasper S. Phelps, Minsu Kim, Helen H. Yang, Arie Matsliah, Zaki Ajabi, Eric Perlman, Kevin M. Delgado, Mohammed Abdal Monium Osman, Christopher K. Salmon, Jay Gager, Benjamin Silverman, Sophia Renauld, Farzaan Salman, Janki Patel, Matthew F. Collie, Jingxuan Fan, Diego A. Pacheco, Yunzhi Zhao, Wenyi Zhang, Laia Serratosa Capdevila, Ruairí J.V. Roberts, Eva J. Munnelly, Nina Griggs, Helen Langley, Borja Moya-Llamas, Zuoyu Zhang, Ryan T. Maloney, Szi-chieh Yu, Amy R. Sterling, Marissa Sorek, Krzysztof Kruk, Nikitas Serafetinidis, Serene Dhawan, Finja Klemm, Paul Brooks, Ellen Lesser, Jessica M. Jones, Sara E. Pierce-Lundgren, Su-Yee Lee, Yichen Luo, Andrew P. Cook, Theresa H. McKim, Dimitrios Stasi Giakoumas, Benjamin Gorko, Emily C. Kophs, Tjalda Falt, Alexa M. Negron-Morales, Austin Burke, James Hebditch, Kyle P. Willie, Ryan Willie, Sergiy Popovych, Nico Kemnitz, Dodam Ih, Kisuk Lee, Ran Lu, Akhilesh Halageri, J. Alexander Bae, Ben Jourdan, Gregory Schwartzman, Damian D. Demarest, Emily Behnke, Doug Bland, Anne Kristiansen, Jaime Skelton, Tom Stocks, Dustin Garner, Anthony Hernandez, Sandeep Kumar, The BANC-FlyWire Consortium, Kevin C. Daly, Sven Dorkenwald, Forrest Collman, Marie P. Suver, Lisa M. Fenk, Michael J. Pankratz, Zepeng Yao, Stephen J. Huston, Tomke Stürner, Gregory S.X.E. Jefferis, Katharina Eichler, Andrew M. Seeds, Stefanie Hampel, Sweta Agrawal, Tatsuo S. Okubo, Meet Zandawala, Thomas Macrina, Diane-Yayra Adjavon, Jan Funke, John C. Tuthill, Anthony Azevedo, H. Sebastian Seung, Benjamin L. de Bivort, Mala Murthy, Jan Drugowitsch, Rachel I. Wilson, Wei-Chung Allen Lee

## Abstract

Just as genomes revolutionized molecular genetics, connectomes (maps of neurons and synapses) are transforming neuroscience. To date, the only species with complete connectomes are worms^1–3^ and sea squirts^4^ (10^3^-10^4^ synapses). By contrast, the fruit fly is more complex (10^8^ synaptic connections), with a brain that supports learning and spatial memory^5,6^ and an intricate ventral nerve cord analogous to the vertebrate spinal cord^7–11^. Here we report the first densely reconstructed adult fly connectome that unites the brain and ventral nerve cord, and we leverage this resource to investigate principles of neural control. We show that effector neurons (motor neurons, endocrine cells and efferent neurons targeting the viscera) are primarily influenced by sensory neurons in the same body part, forming local feedback loops. These local loops are linked by long-range circuits involving ascending and descending neurons organized into behavior-centric modules. Single ascending and descending neurons are often positioned to influence the voluntary movements of multiple body parts, together with the endocrine cells or visceral organs that support those movements. Brain regions involved in learning and navigation supervise these circuits. These results reveal an architecture that is distributed, parallelized and embodied, reminiscent of distributed control architectures in engineered systems^12,13^.

## Main

A coherent understanding of the embodied nervous system remains a central challenge of neurobiology. The fruit fly *Drosophila melanogaster* is the most complex organism for which this milestone is currently within reach. Recent work has yielded connectomes for the adult *Drosophila* brain^14–17^ and ventral nerve cord (VNC)^7–11^. These structures are analogous to the brain and spinal cord of vertebrates, but they contain fewer neurons, making them tractable for complete connectomes (brain: ∼140,000 neurons, VNC: ∼20,000 neurons). The fly brain and VNC are connected by ∼1300 descending neurons (DNs)^18–21^, which have their cell bodies in the brain and project to the VNC, and ∼1900 ascending neurons (ANs)^21–25^. However, the existing fly brain^14–17^ and VNC^7–11^ connectomes were collected separately, and so DNs and ANs are fragmentary in these datasets, though cross-mapping of some cell types have allowed some ‘bridging’ analyses^21^. A unified *Drosophila* connectome would allow us to trace the pathways that connect the brain, VNC and body.

Such a connectome would also shed light on the architecture of behavioral control. Different regions of the central nervous system (CNS) have specialized functions—and this is true in arthropods just as in vertebrates^26^—but we lack a detailed understanding of the overall control architecture in any complex neural system. In principle, behavioral control might flow through a central pathway for perception, action selection and motor coordination; alternatively, it might be decentralized and distributed across many feedback control modules that are loosely coupled in a hierarchical manner. These alternative scenarios are debated in the literature on vertebrate intelligence, insect intelligence and artificial intelligence^13,27–34^. A unified adult *Drosophila* connectome would place important constraints on this debate. Adult flies solve many of the basic control problems that confront other limbed species, including vertebrates^30^.

In this study, we describe the first unified and embodied brain-and-cord connectome of an adult fly. To analyze this dataset, we develop an ‘influence’ network metric to estimate the magnitude of the functional connection between any pair of cells, and we apply this at scale to the entire nervous system. We show that the strongest influences on effector neurons are generally local sensory signals, forming a distributed set of tight feedback loops. Long-range connections involving ANs and DNs coordinate these local loops. Many of these AN/DN circuits can be linked to specific behaviors, such as escape, feeding, reproduction and locomotion. We describe the interactions between these circuits, and we explicitly link these circuits to supervisory brain regions involved in learning and navigation. Our results establish clear empirical support for theories of behavioral control organized around distributed sensory-motor modules, where ‘cognitive’ regions are supervisory but not essential for action.

## Results

### An open-source brain-and-nerve-cord connectome

We generated a serial-section electron microscopy (EM) volume of the connected brain and nerve cord from an adult female *D. melanogaster* at synapse resolution (4×4×45 nm^3^) (**Fig. 1a**). Using our semi-automated sectioning and imaging platform (GridTape7) (**Extended Data Fig. 1a**), we collected 7,010 serial sections onto film-coated tape compatible with transmission EM. This approach enabled visualization of fine neural processes (<200 nm), synaptic vesicles (∼40 nm) and synaptic clefts (∼10-20 nm). After imaging each section, we computationally reassembled the entire Brain and Nerve Cord dataset (BANC, pronounced “bank”) into a 3D volume^8,35^. We then used convolutional neural networks (CNNs) to automatically segment and reconstruct individual cells^8,35^, nuclei and mitochondria (**Fig. 1b**). To proofread and annotate the expected ∼160,000 neurons^10,15^ in the dataset, we followed the approach created by FlyWire for the whole-brain connectome (FAFB-FlyWire)^15,36,37^. We used automatically identified nuclei to account for all neurons with their cell bodies in the CNS. For neurons with cell bodies outside the CNS (e.g., sensory neurons), we manually identified 48 nerves^38–41^ and verified that each axon in these nerves was associated with a segmented neuron. For neurons traversing the neck connective, we verified that every axon at both anterior and posterior neck levels was associated with a segmented neuron. Both the lamina and the ocellar ganglion are missing (see Methods), deficits shared with the emerging MaleCNS project^42^. These regions are present in FAFB, a prior full adult fly brain dataset^16^. A team of 155 proofreaders corrected errors in the automatic segmentation over about 2 years, a total effort of ∼30 work-years (**Fig. 1c**).

**Figure 1:**
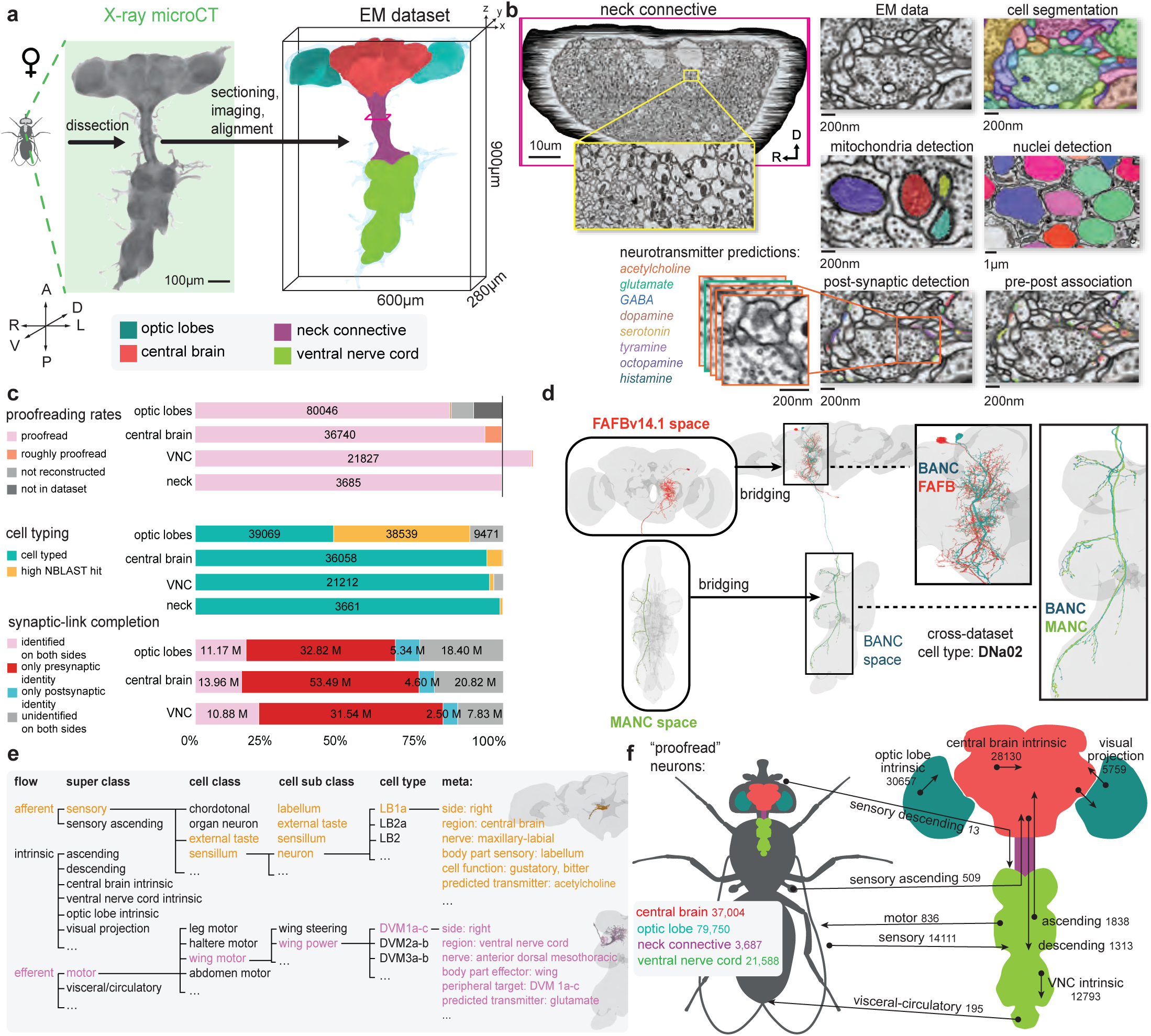
An open-source brain-and-nerve-cord connectome. a. (*left)* X-ray micro-computed tomography (microCT) projection of the BANC sample following dissection, staining, and embedding. (*right*) Rendering of the sample with regions colored. The sample is slightly twisted, but this is corrected using warping registrations (see Methods). A: anterior, P: posterior, D: dorsal, V: ventral, L: left, R: right. b. (*top left*) Aligned EM micrographs through a cross-section of the neck connective ([y=92500](https://ng.banc.community/2025a/neck-connective)) (magenta box in (a)). D: dorsal, R: right. (*yellow box*) Zoom-in of the EM data. (*columns to right*) Example EM image data. Neurons^35,84^, mitochondria ([x: 137533, y: 35220, z: 2493](https://ng.banc.community/2025a/example-mitochondria)), nuclei^85^ ([x: 192977, y: 51679, z: 2493](https://ng.banc.community/2025a/example-nuclei)), postsynaptic locations^50^ ([x: 140988, y :36705, z: 2498](https://ng.banc.community/2025a/example-synapse)) and presynaptic locations (end of yellow lines) were automatically assigned using CNNs. Presynaptic sites were classified by fast-acting neurotransmitter and averaged per-neuron to give neuron-level predictions (**Extended Data Fig. 3**). c. (*top*) Fraction of neurons in divisions of the CNS that have been proofread, based on expected cell counts from other datasets (black line, FAFB counts for brain, MANC counts for the VNC, neck connective counts from ^21^); in the optic lobes, cell types R1-6 and Lai (dark grey) are missing from BANC, indicated by ‘not in dataset’. ‘Proofread’ neurons are manually verified to contain all expected arbors (i.e. axon, soma, dendrite for intrinsic neurons), ‘roughly proofread’ neurons have a soma/primary neurite tract but have some medium to large missing arbor, see Methods. Higher VNC counts for BANC vs. MANC is mainly due to higher recovery of sensory and sensory ascending neurons, which were frequently damaged in MANC. (*middle*) Fraction of proofread neurons in the BANC matched with a cell type in another connectome. ‘Cell typed’ means matched by experts based on morphology and/or connectivity matching; ‘high NBLaSt hit’^44^ means an automated assignment not yet evaluated by an expert. (*bottom*) Fraction of synapses with ‘identified’ (147622 neurons, proofread + roughly proofread + neuronal fragments large enough to be cell typed) pre/post partners. CNS inventory inferred from summing counts from FAFB and MANC. Completion percentages are for BANC v821, but subsequent analyses are performed in v626. d. Neurons were matched to metadata from previous projects by transforming morphologies from other connectomes^8,10,11,14,15,17^ into BANC space^83^. We used NBLAST^44^ to identify morphological matches. An example using DNa02 is shown. [Neuroglancer link](https://ng.banc.community/2025a/DNa02) for morphology, [Codex link](https://codex.flywire.ai/app/search?dataset=banc&filter_string=cell_type+%3D%3D+DNa02&sort_by=&page_size=100) for metadata/connectivity. e. Hierarchy of cell annotations, following previous work^17,86^, but with clearer terms. For example: LB1a ([Neuroglancer link](https://ng.banc.community/2025a/LB1a), [Codex search](https://codex.flywire.ai/app/search?dataset=banc&filter_string=cell_type+%3D%3D+LB1a&sort_by=&page_size=100)) and DVm1a-c ([Neuroglancer link](https://ng.banc.community/2025a/DVM1ac), [Codex search](https://codex.flywire.ai/app/search?dataset=banc&filter_string=cell_type+%3D%3D+DVM1a-c&sort_by=&page_size=100)). See our metadata categories and terms (**Supplementary Data 1**). f. Proofread neurons (totaling 142719) in the BANC by region and super class, for v821.

We assigned cell type labels by automatically identifying potential matches between BANC neurons and earlier datasets^8–10,14,15,17,43^ based on neuron morphology and position (using NBLAST^44^, **Fig. 1c-e**, **Extended Data Fig. 1e**) and based on connectivity (using NTAC^45^). We then manually reviewed and corrected these cell type matches; this process is largely fulfilled but is still ongoing in the optic lobes (**Fig. 1c**). Some neurons are still not cross-matched (10% of BANC neurons excluding the optic lobes), and some of these neurons likely cannot be matched even with more effort, due to inter-dataset variability in cell morphology^46–48^. Inter-dataset variability can result from genetic variation, developmental noise and limitations in data quality or reconstruction. Importantly, in the course of making cell type assignments, we generated the first comprehensive accounting of DN and AN cell types across the neck connective (**Extended Data Fig. 2a**), and we matched AN/DN cell-type labels across the existing whole-brain connectome^15^ and VNC connectomes^8–10^.

To automatically identify synapses in the BANC, we trained another CNN^49,50^ to predict pre- and postsynaptic locations with high accuracy (F-score: 0.83, precision: 0.87, recall: 0.78; **Extended Data Fig. 1b**). Overall, 72% of detected presynaptic links and 23% of postsynaptic links have been attached to identified (proofread and/or cell-typed) cells; 17% of synapses have identified cells on both sides (**Fig. 1c**, **Extended Data Fig. 1c,d**). Comparing the normalized synaptic count between all pairs of cross-matched, identified cell types in the CNS revealed strong concordance between the BANC and other adult fly connectomes (**Extended Data Fig. 1f**), and medium to strong connections present in FAFB and MANC are commonly present in BANC as well (**Extended Data Fig. 1g**).

We used another CNN to predict the neurotransmitter released by each neuron^51^ (**Extended Data Fig. 3a-d**). Our identifications of neurons releasing acetylcholine, glutamate, GABA, dopamine, serotonin and octopamine largely agree with previous predictions^11,51^ (**Extended Data Fig. 3e**). We also used this approach to identify cells that release tyramine and histamine, which have not been previously incorporated into automatic neurotransmitter predictions (**Extended Data Fig. 3a-d**).

Next, we identified many cell types linking the CNS with the rest of the body (**Fig. 1c-f**). To do so, we annotated BANC cells based on literature review, neuron matching and refined labels from prior connectomes. For example, we identified motor neurons targeting muscles of the legs, wings, halteres, antennae, eyes, neck, crop, pharynx, proboscis, salivary glands and uterus^8,10,52–59^. We found putative sensory nociceptors from the abdomen^10,60^, sensory neurons from the aorta^61^, the cibarium^62^ (the pre-oral food chamber), putative oxygen-sensing neurons in the abdomen^63,64^ and sensory neurons from the abdominal terminalia^57,65^. We identified multiple distinct types of endocrine cells in the brain and VNC, many of which could be matched with the neuropeptides they release^64,66–70^ and their sites of action, including the ureter^71,72^, neurohemal release sites, the digestive tract^61,73^, and the reproductive tract^74–76^. We also identified chemosensory, tactile and proprioceptive afferents from the head, eyes, antennae, proboscis, legs, abdomen, wings and halteres^10,41,77–81^. Taken together, these cell type identifications make the BANC a highly ‘embodied’ connectome, with explicit connections to specific muscles, sense organs and viscera throughout the body.

Our ability to describe all these connections relied crucially on BANC being an open science effort^36^ since July 2023, and this project continues to grow with ongoing community input. Users can visualize an archived version of our image data on BossDB (https://doi.org/10.60533/boss-2025-941r), as well as the latest version via Neuroglancer^82^. Users can add annotations through CAVE^37^, and they can also browse metadata and connectivity data via FlyWire Codex^15^ (codex.flywire.ai/banc) and CAVE^37^, as well as programmatically^83^. Neuron morphologies, synapse detections, annotations, indirect connectivity scores, analysis code and more are available as direct downloads from the Harvard Dataverse, a DOI-minting repository (https://doi.org/10.7910/DVN/8TFGGB) (**Extended Data Fig. 1h**). We have also modified typology annotations (super class, cell class, cell sub class, cell function, body part) for the whole female brain (FAFB) and whole male VNC (MANC) connectomes^10,11,15–17^ to facilitate comparisons between these datasets and our work in BANC (**Supplementary Data 1-3**).

### A metric of influence

To interpret a whole brain-and-cord connectome, we need a way to estimate the influence of neuron A on neuron B, for any pair of neurons. To date, there has been no computationally efficient method of estimating these influences. Efficiency is crucial, as there are billions of pairwise interactions between neurons in the full CNS. It would be ideal to precompute all these influences, so that users can simply query any neuron pair of interest.

To tackle this problem, we developed an approach based on a linear dynamical systems description of signal propagation^87–90^ across the brain and ventral nerve cord. Specifically, to compute the influence of one or more source neurons on any target neuron(s), we determine the effect of injecting a sustained signal into the source neurons, taking every downstream neuron’s ‘activation’ as the weighted sum of its inputs. The weight is the number of synapses in that input connection^91^, as a fraction of the postsynaptic cell’s total synaptic input. For a target cell of interest, we take its steady-state response (**Fig. 2a**), log-transform it, and add a constant to ensure that the result is nonnegative. The metric (called ‘adjusted influence’) is inversely proportional to network distance from source to target (**Fig. 2b**, **Extended Data Fig. 4a**). Indeed, adjusted influence is in close agreement with previous network distance metrics^15,22,46^, and like previous distance metrics^15,22,46^, adjusted influence is an unsigned quantity. However, unlike those metrics, our metric is deterministic, linear and computationally efficient. For example, our metric does not involve manually assigning ‘layers’ of neurons for expensive connectivity matrix multiplications (unlike the ‘effective connectivity’ metric)^6^. In essence, the adjusted influence metric is simply an efficient way to estimate the effective distance between any pair of cells, and because it is efficient, it is straightforward to scale it up to quantify all pairwise distances in the connectome. This allowed us to precompute the pairwise adjusted influence of all individual neurons across the brain-and-cord onto all other individual neurons, yielding ∼20 billion scores in total. Across the BANC, the modal pairwise adjusted influence score is 14 for direct connections and 8 for indirect connections (**Fig. 2c**). All scores are available to users via codex.flywire.ai/banc; we also provide code that allows users to compute these scores on demand^92^.

**Figure 2:**
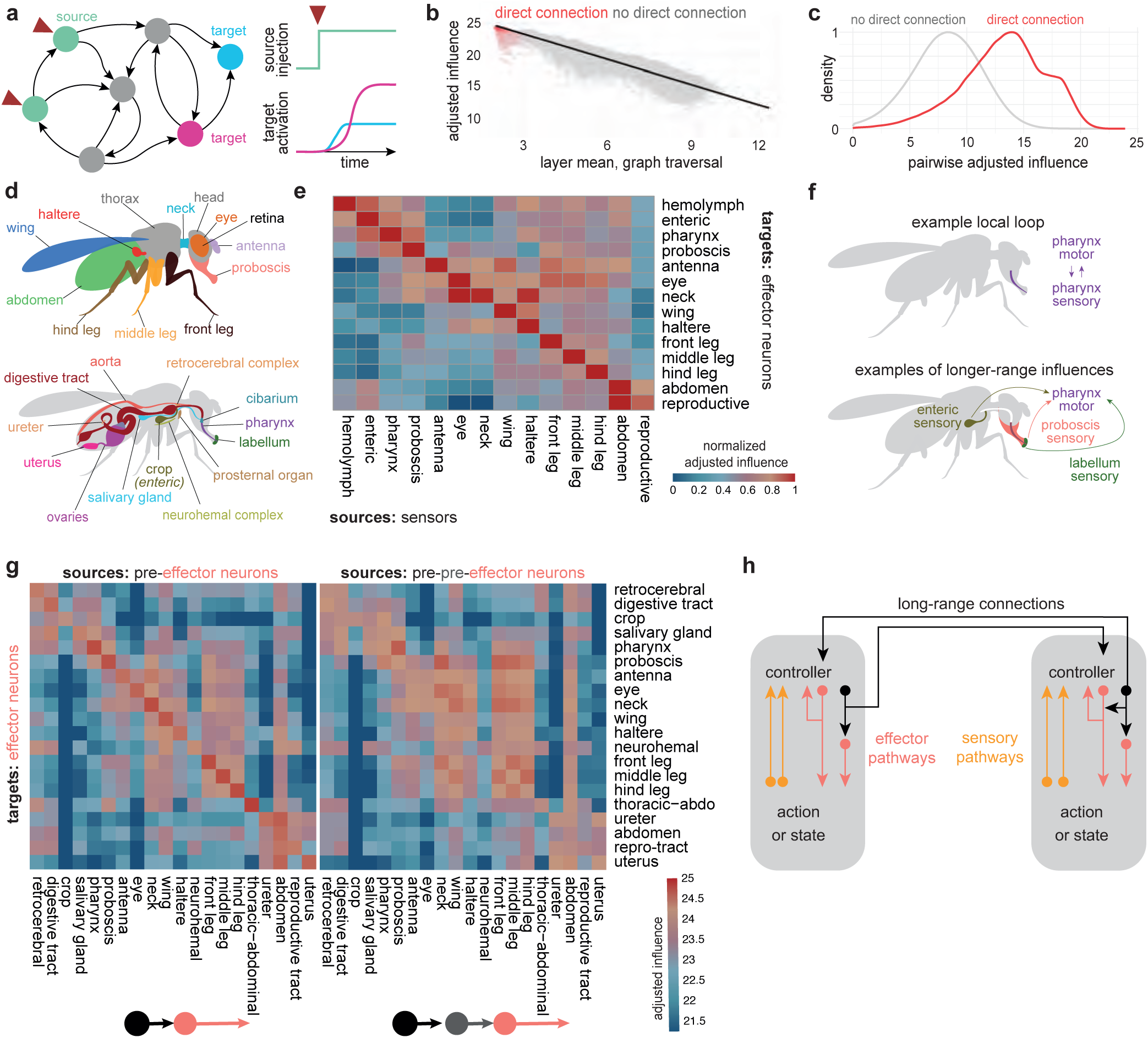
Linking sensors and effectors through local and long-range circuits. a. Influence of source cells on target cells is estimated via linear dynamical modeling. b. Adjusted influence is proportional to the number of network ‘layers’ in a graph traversal model^46^, as expected (see Methods). Adjusted influence is the log(influence), plus a constant (to ensure non-negativity). Direct and indirect connections are shown in red and gray, respectively. Here the source neurons are olfactory receptor neurons in the FAFB dataset^17^, and adjusted influence is averaged over neurons in the source and target groups. Regression line in black (R^2^=0.94, n = 94278 neuron pairs). Note that strong influence can come from either direct or indirect connectivity. c. Distribution of adjusted influence scores between all ANs (1839) and DNs (1313) and all other neurons (155936). Direct and indirect connections are shown in separate histograms, with the peak of each histogram normalized to its own maximum. d. Body parts associated with effector neurons. Not all neurohemal organs shown. [Neuroglancer link](https://ng.banc.community/2025a/efferent-body-parts), explore on [Codex here](https://codex.flywire.ai/app/body_parts?dataset=banc). e. Adjusted influence of sensory neurons (columns) onto effector neurons. Influence is summed over body part before applying Eq. 9 (see Methods). Each row is minmax normalized to the same range (0-1), to make clear which sensor groups are most influential for each effector group (14410 sensory neurons, 1031 effector neurons). ‘Hemolymph’ includes aorta, neurohemal, and retrocerebral complexes; ‘enteric’ includes digestive tract, crop, salivary glands; ‘reproductive’ includes uterus and ovaries; ‘neck’ includes prosternal organ; ‘pharynx’ includes pharynx and cibarium. Putative sensory neurons that could not be matched to body parts are omitted (3188), as are photoreceptors of the retina and ocelli, which are only partially contained in the dataset. Adjusted influence is significantly higher within a body part than across body parts, Wilcoxon rank-sum test, one-sided: within (14 values) > across (182 values), p=4.53×10^-10^. f. Schematic: an example local loop (top) that is also linked to specific sensors via long-range connections (bottom). Connections from sensory neurons to effector neurons are generally indirect, mediated by intrinsic cells of the brain or VNC. g. Adjusted influence of pre-effector neurons on effector neurons (left) and pre-pre-effectors on effector neurons (right). A pre-effector is a neuron directly presynaptic to an effector neuron; a pre-pre effector is defined analogously. Data are for 1031 effector neurons, 38355 pre-effectors, 62389 pre-pre effectors. Two-way ANOVA (Type-III SS) found significant source×target interactions for both, p < 1×10^-15^. h. Schematic: Sensory neurons and effector neurons are connected within functional modules, largely via intrinsic neurons. Effector neuron pathways also influence each other, particularly within a module. Modules are weakly connected via long-range connections. Within each controller, the intrinsic circuitry of the CNS also shapes motor output (e.g., via central pattern generators).

In the following sections, we say A “influences” B, as shorthand for a high adjusted influence score (A→B). These scores do not demonstrate functional connections, and they are no substitute for experiments. The value of these scores is that they allow us to make provisional inferences on a large scale. In the sections that follow, we will use influence scores to make inferences, and to bolster these inferences, we will show example circuit motifs showcasing instances of relatively direct connections. These inferences are merely predictions, and their value is to generate testable hypotheses.

### Modules for local feedback control

Other large-scale connectome analyses have focused mainly on cells deep in the CNS^5,6,93^. Here, we take a complementary approach; we start by focusing on sensors and effectors. A ‘sensor’ is a presumptive peripheral sensory neuron (either external or internal), and an ‘effector neuron’ is defined here as a presumptive motor neuron, endocrine cell, or efferent neuron targeting the viscera, as these are the ‘effectors’ of the nervous system (**Fig. 2d**). Importantly, sensors are distributed across the body, and effector neurons are widely distributed as well: the brain contains motor neurons that control the eyes, antennae, mouth parts, as well as the foregut^94^, while the VNC contains motor neurons that control the legs, wings, halteres, abdomen, reproductive organs and hindgut^7^. Similarly, endocrine cells are found in both the brain and the VNC^95,96^. As an embodied brain-and-VNC connectome, the BANC offers a new opportunity to reconsider sensor-effector relationships.

We found that most groups of effector neurons receive their strongest influence from sensors in the same body part (**Fig. 2e**, **Extended Data Fig. 4b**). For example, we find that pharynx motor neurons are most strongly influenced by pharynx sensory cells. Naturally, pharynx movements will immediately alter the activity of pharynx sensory neurons. Thus, our results imply that pharynx motor neurons and sensory neurons form a reciprocal feedback loop (**Fig. 2f**). We find evidence of similar loops in almost every body part (**Fig. 2e**, **Extended Data Fig. 4b**).

At the same time, the BANC dataset also shows that each local loop is influenced by a select group of more distant sensors in functionally related body parts (**Fig. 2e**, **Extended Data Fig. 4c,d**). For example, pharynx motor neurons are influenced by sensors in the crop, labellum, and proboscis (**Fig. 2e**). These longer-range connections can also be seen as forming feedback loops: for example, pharynx movements during feeding should trigger not only immediate sensory signals in the pharynx, but also more delayed sensory signals in the crop, which might then (for example) limit feeding if the crop is filling too quickly (**Fig. 2f**). In this way, long-range loops can provide important feedback signals that local loops cannot directly access^97^.

In addition, we found that many effector pathways are positioned to influence multiple effectors, and these patterns are also primarily local. Specifically, we identified all the neurons directly presynaptic to any effector neuron (we call these “pre-effectors”); we then computed the influence of each pre-effector onto the entire population of effectors. This analysis showed that pre-effectors tend to influence effectors distributed across multiple body parts, but these influences are selective, and the strongest influences of pre-effectors are their local influences (**Fig. 2g**). We obtained a similar result when stepped back one layer, starting with the neurons directly presynaptic to any pre-effector (“pre-pre-effectors”, **Fig. 2g**). These results argue that effector recruitment is coordinated across body parts, but the strongest effector pathways are local, with weaker coupling at longer spatial scales (**Extended Data Fig. 4e**).

In summary, these analyses argue that CNS output is primarily under the control of local modules. Within each module, sensory neurons influence effector neurons (largely indirectly, via intrinsic neurons); meanwhile, effector neurons drive changes in body state, which produces sensory feedback (**Fig. 2h**). At the same time, within each module, effector pathways also influence each other through CNS circuitry (**Fig. 2h**). Finally, different modules are weakly coupled via longer-range connections. Our findings argue that actions and sensations are tightly coupled, behavioral control is highly parallelized and distributed, and actions are controlled by a combination of sensory reafference and internal expectation signals.

### Linking DNs and ANs to behaviors

Thus far, we have seen evidence for strong local feedback loops, linked by selective longer-range sensor-effector connections. To better understand these long-range connections, we focused on the neurons that link the brain with the VNC, namely DNs and ANs. DNs and ANs participate in many sensor-to-effector connections: we can see this by finding the shortest paths from each sensory neuron to each effector neuron, and then computing the proportion of those paths that contain a given cell of interest. This value is high for DNs and ANs (**Fig. 3a**, **Extended Data Fig. 7a**), which implies that DNs and ANs play key roles in bridging the brain and VNC to connect sensory neurons and with effector neurons.

**Figure 3:**
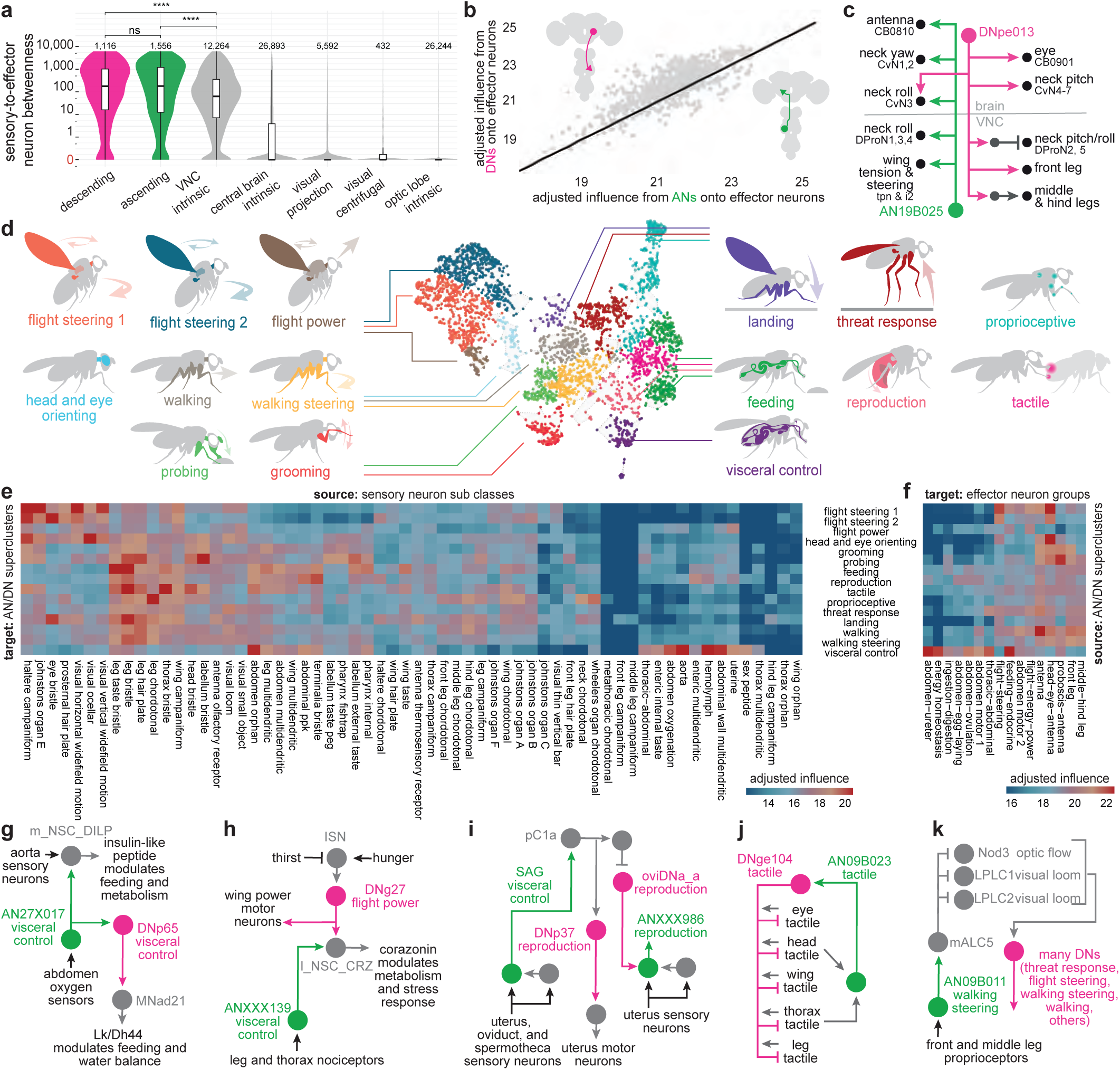
Clustering ANs and DNs into behavior-centric modules. a. “Betweenness” of selected cell super classes, based on the shortest paths from all sensory neurons to all effector neurons. A cell with high betweenness participates in many short paths between sensory neurons and effector neurons. Betweenness varied significantly across super classes (Kruskal-Wallis, p<2.2×10^-16^). Post-hoc Dunn tests with Holm correction (α=0.05) showed that ascending (ANs), descending (DNs) and VNC intrinsic neurons had higher betweenness than all other super classes (all p < 1.5×10“^3^). VNC intrinsic neurons also have high betweenness because the VNC is enriched in sensory and effector neurons (35% of VNC neurons are sensory and 4% are effectors, compared with 16% and 0.5% in the central brain). Pairwise Wilcoxon rank-sum tests with Holm correction (α=0.05) showed significant differences between the medians for ascending vs VNC intrinsic and descending vs VNC intrinsic, but not ascending vs descending (ns p>0.05, **** p<2.09×10^-24^). Boxes show a middle line for the median; box ends = 25th and 75th percentiles of non-logged betweenness. b. Adjusted influence on each effector neuron from DNs versus ANs. Black, unity line. Insets: a DN soma is located in the head, whereas an AN soma is located in the body. c. An example of an AN and DN with strong adjusted influence on effector neurons in multiple body parts. [Neuroglancer link](https://ng.banc.community/2025a/figure-2h), Codex network. d. UMAP embedding of all ANs and DNs based on cosine similarity between their direct connectivity to proofread neurons. Neurons are colored by supercluster membership. [Neuroglancer link](https://ng.banc.community/2025a/an-dn-super-clusters), ANs [here](https://ng.banc.community/2025a/ascending) and DNs [here](https://ng.banc.community/2025a/descending). See **Extended Data Fig. 7**. e. Adjusted influence onto each AN/DN supercluster from select groups of sensory neurons. Superclusters are rows; sensory neurons are columns. A subset of visual project neurons were used to determine processed visual streams from the optic lobes^129^’^143^’^145–158^, see Methods. f. Adjusted influence from each supercluster onto co-controlled groups of effector neurons (**Extended Data Fig. 7**). Superclusters are rows; effector neurons are columns. g. Example circuit involving visceral control ANs and DNs. [Neuroglancer link](https://ng.banc.community/2025a/figure-3g), Codex network. Here and elsewhere, where we show circuit vignettes, it should be noted that we are depicting only a small subset of the inputs and/or outputs of the neurons in question. Our vignettes focus mainly on direct connections, but high influence values often reflect indirect connections. h. Example circuit involving the flight power supercluster and visceral control supercluster. [Neuroglancer link](https://ng.banc.community/2025a/figure-3h), Codex network. i. Example circuit for coordinated visceral sensing and reproductive control. SAG (also called ANXXX983) and ANXXX986 are female-specific^8,21^. [Neuroglancer link](https://ng.banc.community/2025a/figure-3i). Codex network. j. Example circuit involving a DN in the tactile supercluster. [Neuroglancer link](https://ng.banc.community/2025a/figure-3j), Codex network. k. Example circuit involving an AN in the walking-steering supercluster [Neuroglancer link](https://ng.banc.community/2025a/figure-3k), Codex network.

It is sometimes suggested that DNs send motor commands from the brain to the VNC, whereas ANs send sensory signals and predictive motor signals back from the VNC to the brain^24,98^. However, classic work in other insects has shown that some ANs can send signals from the brain to the VNC^99,100^. And in *Drosophila*, more recent work shows that ANs can form output synapses in the VNC^9,10^, while DNs can form output synapses in the brain^21,101^. The BANC dataset shows clearly that both DNs and ANs have substantial output in both the brain and the VNC (**Extended Data Fig. 5a-c**). Moreover, it shows that most effector neurons are influenced by both DNs and ANs (**Fig. 3b**). The majority of individual DNs exert influence over effector neurons in multiple body parts, and the same is true of ANs (**Extended Data Fig. 5d-h**). For example, DNpe013 influences motor neurons for the eyes, neck and legs, whereas AN19B025 influences motor neurons controlling the antennae, neck and wings (**Fig. 3c**). Together, all these observations imply that DNs and ANs work together to coordinate motor patterns and internal organs in different body parts.

### Clustering DNs and ANs into behavior-centric modules

To identify functional divisions among DNs and ANs, we constructed a map of these neurons based on their direct synaptic connections, both pre- and postsynaptic (**Fig. 3d**). DNs and ANs are intermingled in this map because, as it turns out, their connections are often similar (**Extended Data Fig. 6a**). We verified that cells with similar known functions (11% of the total) are frequently colocalized on this map (**Extended Data Fig. 6b,c**; DNs and ANs with known functions are taken from previous work^9,20,21,24,76,101–132^). We then grouped related clusters of DNs and ANs into superclusters, based on the influence they receive from sensory neurons^21^, as well as the influence they exert onto effector neurons (**Extended Data Fig. 6d-g,7,8a**). Based on the sensory and motor influences associated with each supercluster, as well as the functions of known DNs and ANs, we were able to link each supercluster with a putative behavior (**Fig. 3d**).

For example, one supercluster is most likely associated with threat response behaviors. This supercluster contains all the known DNs associated with escape takeoff (**Extended Data Fig. 6b**), as well as many DNs and ANs with unknown functions. As a group, these DNs and ANs are influenced by visual loom detectors, visual small object detectors, and specific mechanoreceptors (**Fig. 3e**). They output to endocrine neurons that regulate internal state, as well as wing and leg motor neurons (**Fig. 3f**). All this is consistent with the idea that these DNs and ANs trigger evasive maneuvers, while also recruiting the energy stores needed to support these maneuvers.

Another supercluster is most likely involved in reproductive behaviors. As a group, these cells are influenced by tactile sensors, taste sensors and nociceptors (**Fig. 3e**). They influence the uterus and reproductive tract, as well as the neurohemal complexes, which release signals into the circulatory system (**Fig. 3f**).

Using a similar process of inference, we linked other superclusters with walking, walking steering, flight steering, flight power, head-and-eye-orienting, grooming, landing, visceral control, feeding and probing (**Fig. 3d**). The term “probing” refers to tactile sampling prior to feeding initiation^133^; we propose that this behavior is mediated by the supercluster receiving strong input from labellar tactile afferents and external taste sensors (**Fig. 3e**) and exerting coordinated influence over the forelegs, proboscis and pharynx (**Fig. 3f**). Meanwhile, we suggest that a distinct supercluster is associated with feeding: this supercluster receives the highest influence from internal taste sensors (**Fig. 3e**), and it has strong influence over the pharynx, crop and salivary glands, as well as endocrine cells targeting the digestive tract (**Fig. 3f**). The influence of the feeding cluster is strongly correlated with the overall influence of pharynx taste and leg taste sensory neurons (**Extended Data Fig. 7b**).

The visceral control supercluster contains ANs and DNs that seem to coordinate endocrine cells in different body parts (**Fig. 3f**). **Fig. 3g** shows an example circuit involving cells from this supercluster. In this circuit, AN27X017 relays signals from putative abdominal oxygen sensors^64^ (Y.L. and J. T., in preparation) directly to brain endocrine cells that release insulin-like peptide (DILP), which regulates feeding^134^; these ANs converge with the projections of aorta sensory neurons^61^. Meanwhile, AN27X017 also synapses onto DNp65, which targets abdominal leukokinin neurons that regulate feeding and diuresis^135^. This circuit might regulate energy and water balance during physical stress.

Any attempt to put DNs and ANs into categories involves some over-simplification, as many of these cells seem to have multiple functions. Consider, for instance, DNg27, in the flight power supercluster (**Fig. 3h**). This DN synapses onto wing power motor neurons, as well as brain endocrine neurons that release corazonin (which mobilizes energy stores^64,136^). Thus, this DN is positioned to increase flight power, while also releasing energy needed to sustain flight. Some of the excitatory drive to DNg27 comes from interoceptive neurons in the brain that are suppressed by thirst^124,137^; this connection may help control flight power based on water balance, because high flight power involves high metabolic demand and thus water loss via respiration^138^. Meanwhile, the same corazonin neurons downstream from DNg27 are postsynaptic to ANXXX139, an AN in the visceral control supercluster that is positioned to relay signals from putative nociceptors. This AN may respond to painful stimuli by recruiting energy reserves, to prepare for struggle or escape. Like many DNs and ANs, these cells are probably multi-functional.

ANs and DNs can sometimes form extended loops. An example involves SAG-ANs (ANXXX983)^117^. The BANC connectome shows that these cells are downstream from sensory neurons in the uterus, oviduct, and spermatheca (**Fig. 3i**), consistent with their known role as monitors of the reproductive tract^117^. SAG-ANs signal to pC1 cells in the female brain^76,117^, which lie upstream from several DNs in the female reproduction supercluster, including oviDNa_a^76^ and DNp37^139^. DNp37 is positioned to regulate uterine motor neurons^57^, whereas oviDNa_a is positioned to modulate ascending sensory signals from the uterus via interposed ANs (**Fig. 3i**). Together, these cells form an extended feedback loop linking uterus sensory signals with uterus motor neurons.

We found two superclusters with particularly strong sensory associations: one is dominated by tactile influence, and the other by proprioceptive influence (**Fig. 3e**). These cells may be involved in whole-body integration of tactile or proprioceptive cues. For example, DNge104 is a cell in the tactile supercluster that is downstream from tactile afferents across the body (**Fig. 3j**) but also upstream from tactile sensors from those same body parts. Because DNge104 is inhibitory, this circuit could produce tactile contrast enhancement. For example, touching the head or thorax is predicted to excite a specific AN that then increases DNge104 activity, thereby suppressing tactile input to the rest of the body. It is interesting that some DNs and ANs are positioned to primarily influence sensory signals, as targeting a sensory signal can be a powerful way to control a behavior: many sensory neurons will carry a feedback signal to one or more loops, and modulating a feedback signal can cause that loop, in essence, to operate with a different setpoint^58,140,141^.

Even in the behavior-centric superclusters, we can find cells positioned to influence sensory processing. For example, AN09B011 in the walking-steering supercluster (**Fig. 3k**) makes a strong direct connection onto a visual centrifugal neuron (mALC5), which is positioned to suppress neurons with ventral visual fields, including visual optic flow detectors (LPLC1^142^, Nod3^143^) and loom detectors (LPLC2^144^). This AN is directly postsynaptic to many types of leg proprioceptors, and so it might function to relay leg movement information to mALC5, allowing this circuit to suppress visual responses to leg movement^145^ or anticipated ventral optic flow from walking.

In summary, we were able to assign a tentative behavioral role to all DNs and ANs, based on their connectivity. These behavioral interpretations are conjectural, but they should be readily testable. Regardless of the exact behavioral function of these cells, however, what is already clear is that many individual DNs and ANs integrate input from multiple sensory streams, and they are positioned to influence many types of effectors. Our results suggest that DNs and ANs are conceptually analogous to the long-range connections that coordinate behavior-centric control modules in robotic design^12,13^. Behavior-centric control modules can be useful because they reduce the need for centralized planning and coordination: complex behaviors can emerge from these modules even without centralized supervision because these modules essentially supervise each other.

### Specialization and coordination among DNs and ANs

Thus far, we have seen that DNs and ANs can be divided into superclusters. Importantly, the cells in these superclusters are not redundant: their inputs and outputs are specialized. As an illustrative example, consider the head-and-eye-orienting supercluster. Different ANs and DNs in this supercluster are influenced by distinct visual or mechanosensory signals, and they influence different combinations of neck and eye motor neurons (**Fig. 4a-c**). Overall, cells in different superclusters have quite different influences, whereas cells within the same supercluster have subtly different influences (**Fig. 4d**, **Extended Data Fig. 8b**).

**Figure 4:**
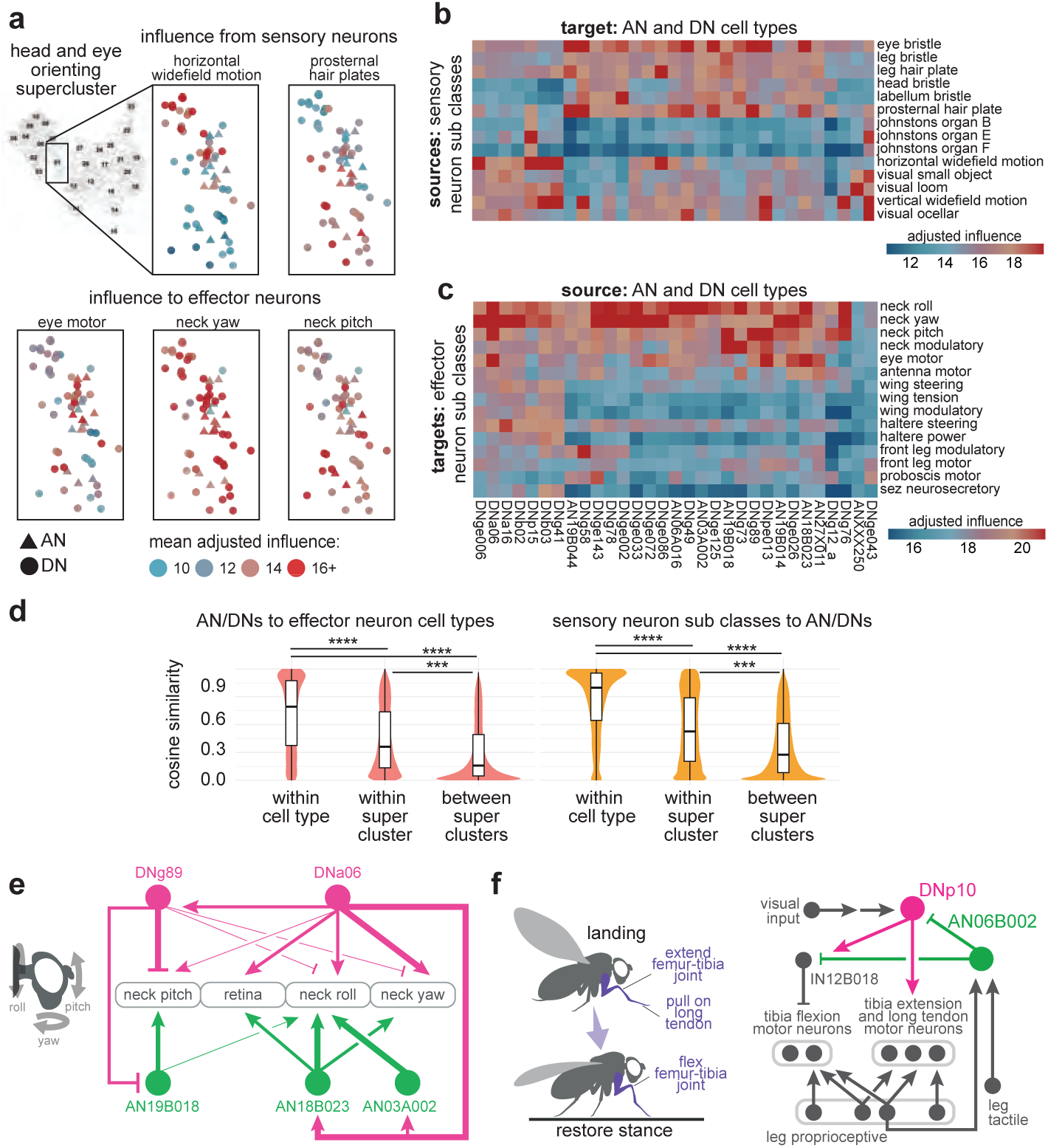
Specializations and coordination within a functional supercluster. a. Enlarged view of the head-and-eye orienting supercluster, taken from the UMAP embedding of all ANs and DNs (Fig. 3d). *Top*: neurons are color-coded by their incoming adjusted influence from two different sensory sources. *Bottom*: neurons are color-coded by their outgoing adjusted influence onto three different effector neuron groups. Each point (AN/DN) is normalized to the same range (0-1), to make clear which sensory neurons are most influential or effector neurons are most influenced [Neuroglancer link](https://ng.banc.community/2025a/head-and-eye-orienting), Codex search. b. Adjusted influence from sensors, for all cell types in the head-and-eye orienting supercluster. Sensory neuron sub classes with adjusted influence over the 95th percentile across all sensory neuron sub classes are included in this plot. c. Adjusted influence onto effector neurons, for these same ANs and DNs. Effector neuron sub classes with adjusted influence from AN/DN types over the 85th percentile across all such influences to effector neurons are included in this plot. d. Cosine similarity of influence scores for ANs/DNs of the same cell type, same supercluster or different superclusters. We compared outgoing effector neuron influences (pink) and incoming sensory neuron influences (yellow). There are significant differences between groups (pairwise Wilcoxon rank-sum tests with Holm correction (α=0.05) (***, p<2×10^-102^, ****, p<1×10^-300^)). e.An example circuit with five cell types in the head and eye orienting supercluster. Thick arrows indicate connections with >100 synapses; intermediate arrows indicate connections with 20-100 synapses; thin arrows indicate connections with 5-20 synapses. This example was chosen to illustrate the concept of diverse but overlapping patterns of connectivity within a supercluster, as well as hierarchical interactions between cells in the same supercluster. [Neuroglancer link](https://ng.banc.community/2025a/figure-4e), Codex network. f. An example circuit with two cell types in the landing supercluster (DNp10^106^, AN06B002). This example was chosen to illustrate the concept that ANs and DNs in the same supercluster can be organized into loops. Note a key role of VNC interneurons in processing sensory feedback. [Neuroglancer link](https://ng.banc.community/2025a/figure-4f), Codex network.

Within a supercluster, cells having different specializations are often linked via direct and/or indirect connections. In some cases, particular DNs or ANs are positioned to recruit (or suppress) many other cells in their home supercluster^101^. Again, the head-and-eye-orienting supercluster provides examples of this. For instance, DNa06 is an excitatory DN with connections onto eye motor neurons as well as neck motor neurons that control all three axes of movement^58^ (roll, pitch, yaw; **Fig. 4c,e**). DNa06 also targets two ANs that are positioned to excite neck and/or eye motor neurons. Meanwhile, DNa06 targets DNg89, which is positioned to inhibit neck-pitch neurons^58^, directly and indirectly through an AN that targets neck-pitch and neck-roll neurons (**Fig. 4c,e**). In short, each DN and AN in this circuit is specialized to influence a specific combination of neck and eye motor neurons, and their interactions might serve to coordinate head and eye movements in different directions.

Within a supercluster, specialized ANs and DNs can also be organized into feedback loops. An example of this from the landing supercluster involves DNp10 and AN06B002. DNp10 drives landing maneuvers in response to looming visual stimuli^106^, and we found this cell is positioned to excite tibial extensor motor neurons and also to inhibit tibial flexor motor neurons via an interposed VNC inhibitory interneuron (**Fig. 4f**), implying that it drives tibia extension during landing. At the same time, we found that AN06B002 is positioned to inhibit DNp10, thereby arresting tibia extension. AN06B002 is postsynaptic to proprioceptive and tactile sensory neurons from the leg (**Fig. 4f**), and so this circuit motif could form a negative feedback loop that arrests tibia extension when the leg has made contact with the surface during landing, allowing the leg to relax into its normal standing posture as the landing maneuver terminates.

In summary, we find that cells in the same supercluster can have specialized connections to sensors and effectors. For each general behavioral task, there is a set of DNs and ANs that link sensors and effectors in diverse, overlapping combinations. Often, these related cells are interconnected, sometimes in loops. These circuits of finely specialized cells should allow for flexible behavioral control that can be rapidly fine-tuned to the current state of the body and the environment.

### Interactions between behavior-centric modules

In a system with behavior-centric modules, there should be ways for one module to influence another. In robotic design, this can help prioritize behaviors, resolve conflicts among behavioral drives, and link related behaviors in sequences^12,13^. Indeed, the BANC dataset reveals a specific pattern of influence among AN/DN superclusters (**Fig. 5a**). Focusing on the strongest of these influences, we can begin to reconstruct relationships between AN/DN behavioral modules (**Fig. 5b**).

**Figure 5:**
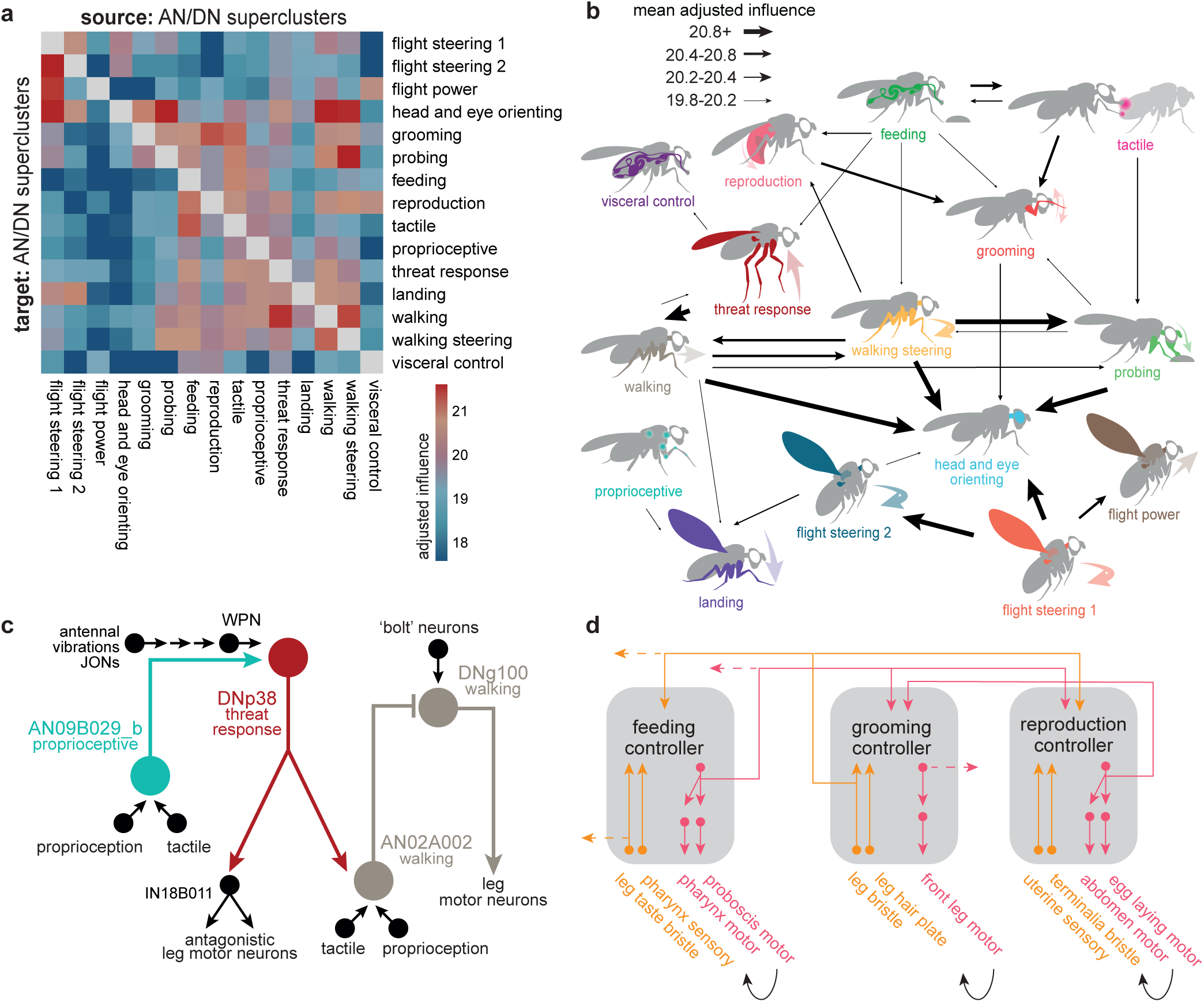
Interactions between behavior-centric modules. a. Adjusted influence of each AN/DN supercluster onto every other supercluster. Values are normalized by the number of cells in each supercluster. Two-way ANOVA (Type-III) on 8572325 observations with 15 sources and targets found a significant source×target interaction (p < 1×10^-15^). Adjusted influence is source size corrected (Eq. 10, see Methods) to control for variation in supercluster size. b. Summary of the strongest adjusted influences between superclusters from panel (a). Here, for clarity, we depict only those influences with an adjusted influence >85th percentile; note that (a) shows all interactions, not just the strongest. Line weights are binned by the adjusted influences shown in (a). c. A circuit illustrating an example of cross-cluster interactions between ANs and DNs. This circuit links neurons in the proprioceptive, threat-response and walking superclusters. Note a key role of VNC interneurons in recruiting leg motor neurons. [Neuroglancer link](https://ng.banc.community/2025a/figure-5c), Codex network. d. Our data are suggestive of an architecture where behavioral modules are linked by selective long-range pathways, creating hierarchical connections among behaviors. Via these connections, one module may recruit or suppress another (subsumption^12^). Here we schematize three conjectured behavioral modules, each constructed primarily from local sensory-motor loops; not all relevant sensors and effectors are shown here. ANs and DNs participate in long-range connections between controllers. Black arrows indicate routes for self-generated sensory feedback (reafference). Dashed lines denote connections to other modules (not shown here). Complex behaviors arise from interactions among low-level feedback loops.

For example, the threat response supercluster strongly influences the walking supercluster (**Fig. 5a,b**), consistent with the idea that threat responses generally require interruption of ongoing walking. Similarly, flight steering and walking steering strongly influence head-and-eye-orienting (**Fig. 5a,b**), reflecting the close coupling between head orientation and steering during flight and walking^159,160^. Finally, walking steering influences probing, a behavior that involves pivoting maneuvers where the fly dances around a food source^161^; this interaction might help coordinate proboscis movements with leg movements.

To better understand the circuits that mediate interactions between superclusters (**Fig. 5a,b**, **Extended Data Fig. 9**), it is useful to drill down to some specific examples. Consider the circuit (**Fig. 5c**) that involves cells from the proprioceptive supercluster (AN09B029_b), the threat response supercluster (DNp38), and the walking supercluster (DNg100 and AN02A002). Here, AN09B029_b sends ascending mechanosensory signals to DNp38, which also receives antennal mechanosensory signals (via WPNs^162^). DNp38 is positioned to drive co-contraction of antagonistic muscle pairs in all the legs, which would likely increase leg stiffness. Thus, this circuit motif might function to integrate whole-body mechanosensory signals to trigger defensive posture stabilization. Meanwhile, DNp38 is also positioned to recruit AN02A002, which inhibits DNg100, a cell in the walking supercluster that is downstream from pro-walking Bolt neurons^105^ and whose direct activation initiates walking^103^. In this manner, a mechanical threat could stabilize the resting stance while also suppressing walking drive.

Overall, the arrangement of influences and direct connections between superclusters (**Fig. 5b**, **Extended Data Fig. 9**) is conceptually analogous to subsumption architecture in robotics (**Fig. 5d**). In such architecture, some behavior-centric modules are positioned to influence, or “subsume”, another module in order to exploit its functionality or override it^12,13^. A set of semi-autonomous modules, loosely linked in a subsumption hierarchy, can produce complex, emergent behaviors^13^.

### Linking behavior-centric modules with other divisions of the nervous system

Finally, we asked how DNs and ANs are integrated with the rest of the brain-and-cord. We began by dividing the CNS into 13 discrete networks, based on each neuron’s direct synaptic connections, using a spectral clustering algorithm that seeks to maximize within-network connectivity while minimizing across-network connectivity (**Fig. 6a**, **Extended Data Fig. 10a**). Our aim was to find large groups of interconnected neurons, as these would be candidate coarse functional divisions of the CNS.

**Figure 6:**
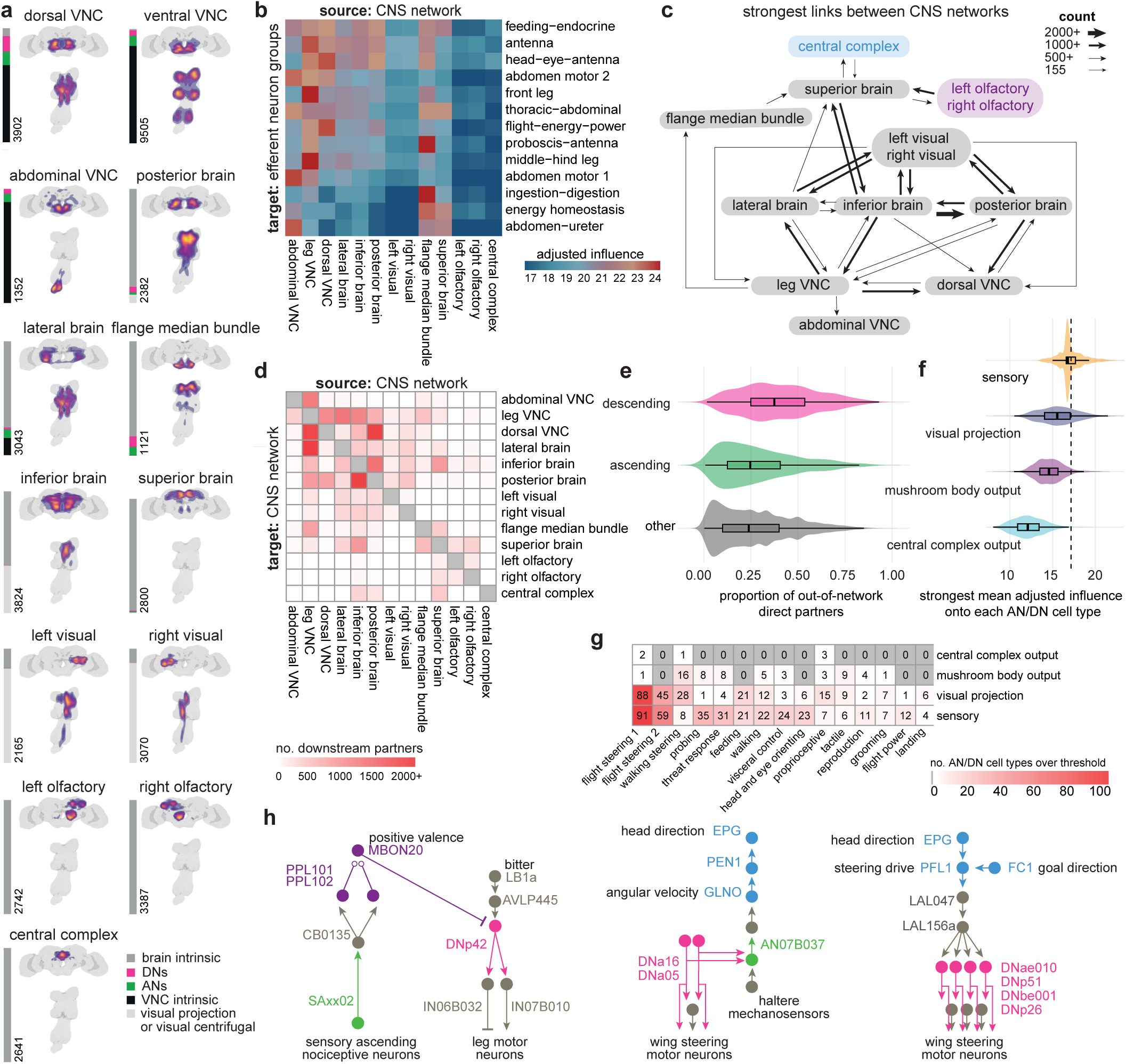
Linking CNS networks with superclusters of ANs and DNs. a. CNS networks, obtained via spectral clustering of 51,502 proofread neurons (excluding sensory, effector and optic lobe intrinsic neurons but including visual projection neurons and visual centrifugal neurons). Each panel includes a 2D kernel density estimation, a stacked bar plot indicating the proportion belonging to each super class, and a cell count. Two pairs of networks are mirror images of each other (olfaction right/left and visual right/left), while all other networks are bilaterally symmetric. Anatomical density images are normalized separately for the brain and VNC, based on a random sample of 100k synapses from each CNS network; the hotter colors indicate a higher density of synapses. b. Adjusted influence of each CNS network on each effector neuron group (effector groups described in **Extended Data Fig. 4**f). Two-way ANOVA (Type-III SS) on 476805 observations with 13 CNS network sources and 13 effector neuron group targets found a significant source×target interaction (p < 1×10^-15^). Most CNS networks (10/13, 77%) have high influence (>median) on at least one effector group. Central complex, right olfactory, and left olfactory networks have zero connections above median (combined probability by chance: p=1.8×10^-12^, binomial test: p=0.046). c. Strongest links between CNS networks. The size of each arrow represents the number of postsynaptic cells in that link. One weaker link is shown (155 cells), because this is the strongest output link of the central complex. d. Link strength between CNS networks, measured as the number of postsynaptic cells in that link. Color scale capped at 2000 cells. e. Out-of-network connections, measured as the proportion of direct partners (up or downstream of the given class with a minimum of 5 synaptic links) each neuron has in another CNS network. “Other” refers to any other neuron in a CNS network that is neither an AN nor DN. ANs and DNs had significantly more out-of-network synaptic partners compared to other (non-AN and non-DN) neurons in their respective CNS networks (p<4.6×10^-6^ for both ANs and DNs, two-sample Kolmogorov-Smirnov test, Holm-corrected (α=0.05), N=20716, K=3 groups, to compare distributions; p<6.2×10^-90^ for DNs and p<3.6×10^-4^ for ANs, Wilcoxon rank-sum test, Holm-corrected (α=0.05), to compare central tendency). f. We took adjusted influence scores from four cell classes (sensory, visual projection, central complex output neuron, and mushroom body output neuron) onto each of the 1018 AN/DN cell types. For each of the four upstream classes, we determined the maximum influence any of its neurons had on each AN/DN cell type; here we show the distributions of these maximum values. The dashed line shows the same, stringent adjusted influence threshold (17.15), developed in **Extended Data Fig. 5**, which is used to threshold cell type selection in (g). Two-sample Kolmogorov–Smirnov tests (Holm-corrected, α=0.05), comparing each group to ‘sensory’ (N=4068 observations; K=4 groups). These tests were significant (Holm-corrected, α=0.05) for visual projection, mushroom body output and central complex output cell types (p < 1×10^-300^). g. Counts for AN/DN cell types above threshold (see f) for each upstream class. Top target cell types as follows. Central complex output: DNa03, DNpe016, DNb01, DNbe006, DNp51, DNa02, DNb09. Mushroom body output: DNge150, DNg104, DNge138, DNge151, DNp42, DNp68, DNge149, DNge152, DNg66, ANXXX977. Visual projection: DNp20, DNp22, DNpe017, DNp18, DNb06, DNp53, DNpe032, ANXXX250, DNg46. h. (*left)* Example circuit connecting mushroom body neurons (purple) to ANs and DNs. Note a key role of VNC interneurons in recruiting leg motor neurons. [Neuroglancer link](https://ng.banc.community/2025a/figure-6g), Codex network. *(middle and right)* Example circuits connecting central complex neurons (blue) to ANs and DNs. Neuroglancer links [here](https://ng.banc.community/2025a/figure-6h-out) and [here](https://ng.banc.community/2025a/figure-6h-out). Codex network.

Notably, most of these CNS networks span the brain and VNC (**Fig. 6a**), and many CNS networks also contain ANs and DNs (**Extended Data Fig. 10b-c**). Most CNS networks also have high influence on at least one group of effector neurons (**Fig. 6b**). Together, these results suggest that behavioral control is highly distributed across CNS networks. The CNS networks with a high influence on effector neurons are directly linked in a dense pattern of reciprocal connectivity (**Fig. 6c,d**, **Extended Data Fig. 10e**). Interestingly, these links are disproportionately composed of DNs: when we counted each neuron’s synaptic partners outside its assigned network, we found DNs had a relatively high proportion of outside partners (**Fig. 6e**). Most AN/DN superclusters are divided between two or three CNS networks (**Extended Data Fig. 10c**), consistent with the notion that DNs often form bridges between networks.

Interestingly, the central complex and the olfactory system emerged as networks with distinctive properties. These networks have relatively low influence on effector neurons (**Fig. 6b**), weak input from other networks (**Fig. 6c,d**), and low AN/DN membership (**Extended Data Fig. 10b**). These networks are likely to have a relatively indirect role in behavioral control: they may merely “supervise” actions, rather than directly controlling actions. This supervision likely involves learning, as the olfactory network includes most of the mushroom body, which is a locus of associative learning, and the central complex is the brain’s navigation center and a locus of spatial learning. Notably, output neurons of the mushroom body and central complex only exert high influence on a small number of ANs and DNs (**Fig. 6f**). Almost all the DNs or ANs with high influence from the central complex are in the walking, walking steering, or flight steering superclusters (**Fig. 6g**, **Extended Data Fig. 10f**). DNs or ANs with high influence from the mushroom body are mainly in the walking steering, threat response, and feeding superclusters (**Fig. 6g**, **Extended Data Fig. 10f**). These results suggest that the central complex and mushroom body supervise specific actions, rather than all actions.

Several example circuits illustrate how these supervisory networks might communicate with lower networks via DNs and ANs (**Fig. 6h**, **Extended Data Fig. 10f,g**). For example, the BANC dataset shows that putative nociceptive cells in the legs (SNaxx02) project directly to the brain, where they are positioned to excite several mushroom body dopamine neurons, including PPL101 and PPL102 (**Fig. 6h**). These dopamine neurons encode negative valence^5,163,164^, and they are positioned to instruct olfactory learning in several mushroom body output neurons, including MBON20^5^. Given the synaptic learning rules governing olfactory learning in the mushroom body, we would expect that these dopamine neurons will “teach” MBON20 to respond selectively to odors lacking negative associations - i.e., odors associated with safety. Notably, MBON20 is positioned to inhibit DNp42, which drives backward walking in response to noxious stimuli^109^. Thus, odors associated with safety should excite MBON20, which is then positioned to suppress avoidance behavior (**Fig. 6h**). This example circuit illustrates how the olfactory network can supervise behavior by interacting with ANs and DNs.

Another example circuit comes from the central complex. In the central complex, angular path integration is driven by an internal estimate of the fly’s rotational velocity, encoded by GLNO neurons^165^. The BANC dataset reveals that GLNO neurons receive a strong disynaptic excitatory input from a specific AN (**Fig. 6h**). This AN receives direct input from DNa16 and DNa05, which likely contribute to steering in flight, via direct and indirect connections onto wing steering motor neurons. Thus, this AN is positioned to send copies of descending flight steering signals back up to the central complex, to update the head direction system in anticipation of an upcoming change in heading. The central complex continuously compares the fly’s estimated head direction against its internal goal direction. This comparison is performed by several cell types, including pfli^6,166,167^, but the DN targets of PFL1 have not been fully identifiable until now, as DNs were fragmentary in available connectomes. The BANC dataset shows that DNs downstream from PFL1 are in fact putative flight steering neurons (**Fig. 6h**). Thus, PFL1 is positioned to compare head direction with its goal direction and to generate corrective steering commands in flight when these directions are misaligned. Again, this example illustrates how the central complex can supervise behavior by interacting with ANs and DNs.

## Discussion

The BANC represents a major advance in scale and complexity compared to other connectomes (*C. elegans*^1,2^*, Ciona intestinalis*^4^, and *Platynereis dumerilii*^3^). Tackling a problem of this scale required us to leverage new methods for semi-automated sectioning and EM imaging, computational section alignment, cell segmentation, synapse identification, neurotransmitter assignment and cell-type matching. Because we could draw on the expertise of a large open science community, we were also able to assemble an embodied connectome with explicit connections to many organ systems.

An embodied connectome of this scale offers new clues about the logic of behavioral control. The “classical” theory is that actions are selected by a centralized executive brain function (sensing→cognition→action, the “classical sandwich”)^168^. The implementation of motor patterns could be delegated to circuits in the spinal cord or insect VNC^30,169,170^ but in the classical view, these motor patterns are selected by descending commands from a centralized executive controller in the brain. Following this classical model, executive functions like action selection and attention have been ascribed to the insect central complex or mushroom body^27,31–34^. A radical alternative is the “ecological” theory, which utterly dispenses with the middle layer of the classical sandwich: it posits direct bidirectional connections between sensing and action (sensing→action), rejecting the need for internal representations^171^. More recently, theories of “embodied cognition” have argued that sensing→action loops could be implemented by a modular, distributed control architecture that is tightly and strategically connected to the body, taking inspiration from robotic design^172^. Theories of embodied cognition do not reject internal representations, but they argue their role is limited — e.g., local efference copy signals might be used to inform locomotor control, or memorized snapshots might be used to guide navigation. Models of embodied cognition predict that most behavioral choices engage distributed networks, consistent with recent data from mammals^173,174^.

Our analysis of a fully embodied connectome provides new insight into how distributed behavioral control might be implemented. First, our results imply that the core elements of behavioral control are a set of local feedback modules, where effector neurons are strongly and specifically influenced by the sensors positioned to monitor the relevant effectors. Local feedback loops are known in the vertebrate spinal cord and brainstem^175,176^, as well as the insect vnc^9,30,170,177–180^, gnathal ganglion^52^, and enteric nervous system^61^. Our study extends prior work to show that local loops are located throughout the CNS and that they are consistently the strongest influences on effector neurons. Second, our analyses draw attention to the ubiquitous recurrent connections between effector pathways, which again are primarily local. These recurrent connections are positioned to alert the CNS to upcoming movements, thereby reducing processing delays and helping to coordinate different effectors. The prominence of local feedback loops and recurrence supports the embodied cognition model of neural function.

At the same time, purposeful behavior also requires long-range coordination among body parts (long-range loops^97^), which are mediated, in part, by ANs and DNs. We found these cells can be divided into superclusters, with each supercluster linking a specific set of sensory cells and effector neurons. We link many superclusters with putative behavioral functions, and we highlight hierarchical connections among superclusters^21,101^. This control architecture — typified by behaviorally specialized long-range connections arranged in a hierarchical manner — is reminiscent of subsumption architecture in robotic design^12,13^, which shares many tenets with the theory of embodied cognition^172^. In these robots, behavioral control is divided among many feedback control modules, each devoted to a behavioral task or aptitude, and high-level modules have the ability to recruit or suppress lower-level modules via hierarchical connections. This type of architecture can also potentially account for some hierarchical relationships among animal behaviors^181,182^.

According to the classical theory, descending signals are merely action commands, which in principle might be packaged into a small number of descending axons^19,21^. To the contrary, in the BANC dataset, the sheer number of DNs (∼1300 cells) is larger than the number of effector neurons (∼1000), and ANs, including sensory ANs, are even more numerous (∼2400). Indeed, within each AN/DN supercluster, we find many fine-grained variations on the same connection pattern, forming parallel pathways with slightly different inputs and/or outputs. This arrangement should promote flexibility, by offering many available action patterns. It should also promote precision, by pre-selecting the specialized action patterns that can result from particular patterns of sensory input. These specializations could explain why, for example, different threat response DNs can produce different escape takeoff maneuvers^183^, and why different steering DNs can produce distinct changes in leg movement^111^. Importantly, DNs and ANs typically influence effector neurons indirectly, via interneurons, and the circuitry of these interneurons often plays a key role in structuring behavioral dynamics^184–187^.

Finally, considering the network structure of the entire brain-and-cord, we found that many CNS networks have a high influence on effector neurons, further supporting the idea that behavioral control is distributed, rather than centralized. A few CNS networks have notably low influence on effector neurons; this is particularly true of the central complex and the olfactory network (which includes most of the mushroom body). Thus, the central complex and mushroom body are unlikely to perform central control of action selection. In addition, we found that the central complex and mushroom body have high influence on only a small subset of DNs and ANs, consistent with previous work showing that major central complex or mushroom body disruptions have quite specific effects on behavior^188,189^. Together, these findings imply that the central complex and mushroom body have a supervisory role and that they supervise a subset of behaviors, not all behaviors. This conclusion is compatible with the idea of embodied cognition (which posits a limited role for memories and other internal representations); it is also compatible with subsumption architecture, where high-level modules supervise low-level modules, but high-level modules are inessential for many behaviors^12,13^.

This project illustrates how insight can arise from new technologies, combined with the accumulation of many small biological facts. Just as early cartographers amalgamated the work of multiple map-makers, we have deliberately amalgamated typology and metadata from prior *Drosophila* connectomes. We anticipate future work will amalgamate the BANC with additional datasets, including the recently released MaleCNS connectome^42^, enabling studies on sex-specific wiring and inter-individual variability. The BANC has been an open science project since 2023 and, as a living public dataset, it will progressively improve as long as users continue to interact with it. This open science effort should generate even more testable experimental hypotheses and, ultimately, new theories.

## Acknowledgements

We thank Aaron Kuan for discussions, advice and assistance with specimen preparation and the imaging pipeline; Philipp Schlegel and Casey Schnieder-Mizell for advice and assistance with computational infrastructure and maintaining open source connectomics tools; members of SixEleven (Jenny Esteban Abelo, Rimalyn Enopli Silay, Anthony Ralph Piccio, Renzo D. Requintina, Micah Colinares Molinas, Shaina Rose Araneta, Jazel R. Digamon, Carine E. Estrella, Marlon Kedet Roda, Michael Manayaga Mates, David Benjamin D. Conquilla, Romilyn Bardonido, Alvin Josh Mandahay, Rey Mark Asares, Brion Glee Baliña Gloria, Aleanne May Hopkins, Jessa Calambro, Christine Joy Maquilan Tac-an, Cj Christine Cordova Tan, Cj Pilapil, Lea Lovely Joy Jimenez Pelingon, Mary Rose Golez, Prince Lloyd Sy Cagampang, Jacquilyn Laude, Ehreca Mae Jongco Pojio, Jayanne Biodes Mayormita, Mike Quibo Tambuco, Madiemie Inguito, Cedrick Earl Bernasol, Roden Rey Nuiz Dizon, Francis Rainer Lindo, Noroddin Balambag, Cathy Pilapil, Tutor, Larben Milagrosa, Clyde Albero, Cesar Jun M. Subtenente, Elison Rovic Monte, Sherwin Salem, Estephani A. Lim, Enriq Gerald Pacaldo, Juliane Maquiling Bulaclac, Yna Bustria Jallorina, Catherine Gasi Colminas, Michael Carlo M. Pabroquez, Jonalyn S. Revilleza, Nina Mae Dulaca, Marchan Manaytay, Rommel Roa, Glenda Malda Dawa, Baby Jane Getizo, April Grace Delantes Alagaban, Krystel Joy Domingo, Mark Lester Avila, Nelsie Panes, Dwight Jumaag, Hugh Ven Kyle Tecson, Jansen Seguido, James Nicle Cano, John David Asis, Althea Lou Alojipan Aligria, Philip Ampo, Mary Grace Ybanez, Charmelyn Urboda, Vanessa C. Dal, Rhosly Jay Bumaya, Regine Salem, Jan Marinie Ferraren, Mark Lester Caño, Joseph Enoc Roilo, Aljon Lastimoso, Romelita Maignos, Kendrick Joules Vinson, Hella Jones Dulay, Celso Gonzales, Raffy Acosta, Irish Quijada Bonso, Johnclyde Saguimpa, Wilbur Hernando De la Torre, Mark Lloyd Pielago, Clerk Derecho, Jaypee Balco, Remer Tancontian, Allien Mae Gogo, Joshua Bañez, Nashra Hadjerol, Jeslie Labaya Bongcales, Chereb Martinez, Ianloyd B. Polinar, Jeanity Dominguez, Charlote B. Solana, Queendolyn Pama, John Niño M. Ugdang, Melcris Damasco, Marianne Marie Villarba, Edrian R. Navaja, Janice Salocot, Daril Bautista, James Andrew A. Campos, Elanur Jimlani Acmad, Jet Dolorosa, Zairene Lenizo, Rey Adrian Candilada, Logic Domme Sauro Ybañez, Mark Clint Encallado and Jovie Cano) for proofreading; May Husseini for project administration; Celia David for creating training materials and helping to manage citizen scientists; the management at SixEleven for coordination and professional proofreader management; Alexander Malakov for assistance with software development; Stephen Holtz for discussions on chordotonal organs; Pablo Reimers for ground truth related to LPsP neurotransmission; Noah Pettit for insights on PFL1 neurons; Lydia Hamburg for discussions on connectome analysis; Anton Miroschnikow for discussions on Hugin neurons; Rosalinda Maggio for unpublished insights into neurohormone expression in VNC neurons; Lisa Marin for discussions on MANC annotations; Yijie Yin and Volker Hartenstein for discussions on hemilineages; Lou Scheffer for comments on our preprint and data products, and discussions on synaptic completion rates across datasets; Richard Schalek for help with X-ray microCT scanning and processing; Yijie Yin for discussion on reporting cell functions; David Masao Zimmerman for advice and discussions on analysis; and Lee and Wilson lab members for feedback on the manuscript. We thank five anonymous reviewers, whose work helped to improve this manuscript. This work was supported by NIH grants R01NS121874 (to B.d.B and W.C.A.L), RF1MH117808 (to J.C.T., H.S.S., and W.C.A.L.), U19NS118246 (to J.D.), U24NS126935 and RF1MH117815 (to M.M. and H.S.S.) to perform large-scale proofreading, annotation, and dissemination of the BANC dataset. Our work has benefited from the O2 High-Performance Compute Cluster, supported by the Research Computing Group, at Harvard Medical School (https://it.hms.harvard.edu/our-services/research-computing). A.S.B. was supported by a Sir Henry Wellcome Postdoctoral Fellowship (222782/Z/21/Z); H.H.Y by a NIH K99 award (K99NS129759); B.d.B. by a Smith Family Odyssey Award; R.M. and B.d.B. by a Harvard/MIT Joint Research Grant; D.A.P by a HHMI Awardee of the Life Sciences Research Foundation Postdoctoral Fellowship (PJ100000343); S.A. by the NIH (R00NS117657); J.C.T by the NIH (R01NS102333 and RF1NS128785) and a New York Stem Cell Foundation Robertson Neuroscience Investigator Award; M.P.S. by the NIH (R01NS140174); M.Z. by the Deutsche Forschungsgemeinschaft (ZA1296/1-1) and NV INBRE (GM103440); K.D. by the NSF (2127379); G.S. by the JSPS (KAKENHI 25K00370) and JST (ASPIRE JPMJAP2302 and CRONOS JPMJCS24K2); M.J.P. by the Deutsche Forschungsgemeinschaft (EXC2151-390873048, PA787/7-3, and PA787/9-3); F.C. by the NIH (UM1NS132253, U24NS139927, RF1MH128840); J.M.J by a HHMI Gilliam Fellowship (GT15790); L.M.F and T.F. by the Max Planck Society; S.D. by the Shanahan Family Foundation; M.A.M.O. by a Kempner Graduate Fellowship; A.M.S. and S.H by the NIH (R01NS121911); G.S.X.E.J. by the NSF (2014862) and MRC (MC_EX_MR/T046279/1); Z.A. by the Alice and Joseph Brooks Fund; S.R. by the NIH (T32GM144273); A.M.N.M by the NIH (R25NS080687); *Z.Y.* by the NIH (R01DK139131); and T.S.O. by the Beijing Natural Science Foundation (IS23084). We thank Bradley Dickerson for his support of S.D. and E.B (NIH grant U01NS131438, NSF grant IOS2221458, Searle Scholar and McKnight Scholar Awards). R.I.W. is an HHMI Investigator.

## The BANC-FlyWire Consortium

Includes the listed authors of this paper and additionally: Anna Verbe^1^, Gabriel A. Nieves-Sanabria^2^, Devon Jones^3^, Zijin Huang^1^, Sofia Pinto^4^, Celia David^3^, Omaris Y De Pablo-Crespo^2^, Emily Ye^5^, Wolf Huetteroth^6^, Zequan Liu^7^, Fernando J. Figueroa Santiago^2^

^1^Princeton Neuroscience Institute, Princeton University,Princeton, NJ, USA

^2^Institute of Neurobiology, University of Puerto RicoMedical Sciences Campus, San Juan, Puerto Rico

^3^Eyewire, Boston, MA, USA

^4^Champalimaud Neuroscience Programme, ChampalimaudCentre for the Unknown, Lisbon, Portugal

^5^Department of Neurobiology, Harvard Medical School,Boston, MA, USA

^6^Department of Biology, Leipzig University, Leipzig, Germany

^7^RWTH Aachen University, Aachen, Germany

## Author Contributions

R.M. and B.D.B. selected and behaviorally characterized the sample fly. M.K., R.J.V.R. and B.M-L. generated synapse ground-truth. J.S.P and M.K. prepared the sample and collected and aligned the EM dataset. S.P., N.K., D.I., K.L., R.L., A.H., A.B. and T.M. at Zetta.ai ran automatic cell, nuclei and mitochondrial segmentation on the dataset as well as synapse prediction. M.K., R.J.V.R. and B.M-L. evaluated the automatic segmentation and synapse prediction. K.M.D., D-Y.A., A.S.B and J.F. collated neurotransmitter ground-truth and developed a fast-acting neurotransmitter prediction for BANC. E.P., S.D. F.C. and A.H. lead data management through CAVE and Neuroglancer. M.M. and H.S.S. managed the FlyWire team and led the proofreading and dissemination efforts. C.S., S.-C.Y. and B.S. oversaw proofreading efforts through SixEleven. Specialist proofreading was conducted by L.S.C., R.J.V.R., H.L., E.M., N.G., B.M-L., A.C., J.G., B.S., A.B., J.H., K.W. and R.W., managed by H.H.Y. FlyWire citizen scientist members K.K. and N.S. contributed proofreading and annotations. A.R.S. and M.S. managed the FlyWire community and onboarded new members. J.S.P., H.H.Y. and A.S.B designed and implemented the annotation metadata scheme for neuronal cell types in BANC. J.S.P. bridged the dataset to template spaces. M.K., H.H.Y. and R.J.V.R. identified and seeded neuronal profiles of the neck and nerves. Key neuron annotations were contributed by H.H.Y., A.S.B., J.S.P., M.K., C.S., F.S., S.D., T.S., F.K., P.B., E.L., S.E.P-L, J.J., S.P., S-Y.L., A.C., B.J., G.S., A.M., D.D, M.J.P., B.G., S.J.H., D.S.G., E.C.K., T.F., M.P.S., L.M.F., Z.Y., K.E., S.H., S.A., T.M., M.Z., Jx.F., D.A.P., J.P., J.T., A.A., J.G., B.S., A.B., J.H., K.W., R.I.W and the BANC-FlyWire Consortium, particularly a large number of motor neurons by A.A. S.E.P-L., Y.L., A.A., J.J., S.A., A.C., M.J.P., D.D., M.Z. and J.T. shared unpublished experimental observations that benefitted this work. The influence metric was designed by Z.A., J.D., A.S.B. and R.I.W. Jx.F. implemented the cascade method for validating the influence metric. J.S.P., E.P., A.M., G.S.X.E.J., K.K., Z.A., Jx.F., H.H.Y., and A.S.B. built programmatic tools for data access and analysis. A.M. disseminated data and analysis results through FlyWire Codex. Data analysis was conducted by A.S.B, M.O., T.S.O., Z.Z. and R.I.W. Circuit vignettes were built by A.S.B, S.R., M.F.C., Jx.F., D.A.P., Y.Z., W.Z, H.H.Y. and R.I.W. A.S.B., W.C.A.L. and R.I.W. wrote the manuscript with feedback from the authors. A.S.B, H.H.Y., J.S.P., M.M., R.I.W. and W.C.A.L. managed the project. M.F.C. and A.R.S. produced graphics and illustrations, H.L. and L.S.C. copy edited the manuscript. Correspondence on cell typing, data and data analysis can go to A.S.B. and J.S.P. Correspondence on FlyWire and proofreading can go to M.M. Correspondence on influence scores can go to J.D. Correspondence on science and data collection can go to R.I.W. and W.C.A.L.

## Methods

### Overview

To produce the Brain and Nerve Cord (BANC), we dissected a CNS sample from an adult female fly, stained and fixed the tissue, sectioned the sample and imaged sections at synapse-resolution. We aligned the image volume into a coherent 3D dataset, and Zetta.ai performed automated image segmentation and synapse detection. SixEleven and Aelysia LTD performed the bulk of the human “proofreading” to fix automated segmentation errors and to identify sufficiently complete neurons. The project’s proofreading and annotation infrastructure was based in Neuroglancer^82^ (proofreading, visualization) and CAVE^37^ (key metadata annotation) and managed together with Yikes LLC. We built custom R and Python tools and code for data analysis and to manage and assign metadata at scale. These tools were built on open source projects including Navis^190^ (Python) and the Natverse^83^ (R). We used established frameworks for metadata assignments where possible, and a small core team of two to four scientists manually reviewed them. The wider research community was openly engaged to improve the BANC, mainly via CAVE^37^ and FlyWire^36^. FlyWire Codex (https://codex.flywire.ai/banc) provides user-friendly access, and reflects continued improvements to the dataset.

### Specimen

The BANC sample came from a female adult fly. We behaviorally screened 5-6 day post-eclosion wild-type *Drosophila melanogaster* (F1 progeny of a w^1118^ × Canton-S cross) female flies^191,192^. The fly used for the BANC dataset turned right 70% of the time over 582 choices when walking in an acrylic Y-maze for 2 hours. We raised the flies on standard cornmeal-dextrose medium at room temperature (∼20 °C) in natural lighting conditions. We collected flies on the day after eclosion, housed them in vials with other flies for 4-5 days, behaviorally tested them and then subsequently housed them individually in vials for ∼1 day at 25°C until dissection.

To dissect the flies, we pinned them individually onto a dissection pad then submerged them in a drop of ice cold Karnovsky’s fixative (2.5% formaldehyde, 2.5% glutaraldehyde in 0.1M cacodylate buffer, pH 7.4) containing 0.04% CaCl2. We removed the legs and proboscis to allow fixative to access the nervous tissue.

Next, we carefully removed the head capsule and the cuticle of the ventral thorax to expose the nervous tissue for dissection. Within 5 minutes, we completely dissected the brain and connected VNC, and we transferred it to an Eppendorf tube containing the same Karnovsky’s fixative. We fixed the sample at 4 °C overnight. On the subsequent day, we washed the sample with 0.02M 3-amino-1,2,4-triazole (A-TRA) in cacodylate buffer (3×10min) and then we stained it with 1% OsO4 in 0.1M A-TRA for 90 minutes on ice. On the same day, we stained the sample with 1% thiocarbohydrazide for 8 minutes at 40 °C, 2% OsO4 (aqueous) at room temperature for 60 minutes, and 1% uranyl acetate in maleate buffer at 4 °C overnight. On the next day, the sample was stained with lead aspartate for 3 hours at 60 °C, then dehydrated in a graded ethanol series, washed with propylene oxide, and infiltrated with 2:1 and 1:2 propylene oxide:LX-112 resin consecutively for 30 minutes each. The sample was then placed in pure LX-112 resin overnight at 4 °C and was embedded in fresh pure resin the following day and cured at 60 °C for 48 hours.

The resin-embedded sample was scanned on a microCT X-ray scanner (Zeiss) before serial sectioning to screen for obvious defects or damage. Importantly, the neck connective appeared intact. The specimen includes the central brain, neck connective, VNC and the medulla, lobula, and lobula plate of the optic lobes. It lacks the lamina (part of the optic lobes), the ocelli, and the ocellar ganglion. Thus, the R1-6 photoreceptors, the lamina intrinsic cell type Lai and the ocellar photoreceptors are missing from BANC (∼10000 cells) and intrinsic neurons that arborize in the lamina and ocellar ganglion are incomplete (cell types: L1, L2, L3, L4, L5, T1, C2, C3, Lat, Lawf1, Lawf2, OCG01, OCG02, OCC01, OCC02, DNp28 and the ocellar local neurons). There is also dissection damage to both the left and right antennal nerves. This has resulted in damage to Johnston’s organ sensory neurons and central brain intrinsic neurons that pass near the nerve entry point, which means that their recovery and cell typing is suboptimal. These defects (of the lamina, ocellar ganglion, and quality of Johnston’s organ neurons) are also true of the maleCNS project, a whole male brain and nerve cord^42^, though BANC’s Johnston’s organ sensory neurons are more complete. The BANC is the only available dataset for which the full female abdominal neuromere is present.

### Serial sectioning

We cut serial 45-50 nm thin sections and collected them on a 7500-slot reel of GridTape (Luxel) as previously described in^7^.

### Transmission electron microscopy (TEM) imaging

We used one TEM (JEOL 1200 EX) with a custom vacuum extension and scintillator (Grant Scientific), 2×2 array of sCMOS cameras (Andor, Zyla 4.2), and custom modified with a reel-to-reel, GridTape imaging stage to acquire the dataset as described previously^7^. Imaging spanned 7.5 calendar months, but 96.5% of the images were acquired during the 4 months of November 2021 to February 2022.

### Missing data

Of the 7010 sections, 6970 (99.43%) were collected and imaged without data loss. Ten (0.14%) have no data due to the section being lost (sections 856, 885, 3755, 5746, 5772, 5778, 5793, 5801,5822 and 5869). Notably none of the losses are consecutive serial sections. One of these losses (3755) was because the section was collected onto the wrong location on the GridTape (not over the slot) and so it could not be imaged with TEM. The other 9 losses were due to the support film rupturing after section collection but before the section could be imaged. An additional 30 sections (0.43%) have partial data: 11 sections are missing all images for the brain: 914, 1462, 5841, 5849, 5888, 5896, 5916, 6207, 6208, 6209 and 6210; 7 sections are missing all images for the VNC: 874, 2784, 2822, 3064, 3102, 4566 and 5840; 12 sections are missing a fraction of brain and/or VNC images: 2828, 2860, 2912, 2986, 3054, 3080, 3586, 3605, 3833, 4648, 4768 and 5935. The large majority of these losses were also caused by partial rupturing of the support film before the tissue was imaged.

### TEM dataset alignment and segmentation

We performed initial BANC image alignment with a custom software pipeline that deployed AlignTK alignment functions (https://mmbios.pitt.edu/aligntk-home) on a computing cluster^7^. We refined the alignment of the data using self-supervised CNNs and online optimization to produce displacement fields that were combined with a global relaxation^193,194^. We next trained a CNN to identify regions that were damaged during serial sectioning.

We then used CNNs to segment the dataset into cells and fragments of cells at 16 x 16 x 45 nm^3^, excluding regions that decreased cell segmentation performance including areas with damage, as well as organelles including nuclei and mitochondria^8,35^. We then ingested the automated segmentation into the Connectome Annotation Versioning Engine (CAVE)^37^ for distributed proofreading.

### Synapse detection

We generated synapses in two-steps: (1) postsynaptic terminal detection and (2) synaptic partner assignment^195^. We pretrained both models with data from FAFB, and we tuned the detection model with an additional 721 (∼21% new) labels from the BANC. The detection operated on 8 x 8 x 45 nm^3^ images, with an output at 16 x 16 x 45 nm^3^. We removed detection objects <3 voxels. Assignment operated at 16 x 16 x 45 nm^3^. We merged terminals with identical assignments that were within 200 nm of each other into a single terminal. This detection is known as synapses_v2 (formerly known as synapses_250226 in our v1-2 preprint) and is available through CAVE. It comprises 218460852 synaptic links, of which 72% of presynaptic ends and 23% of postsynaptic ends are connected to a proofread neuron, i.e. the synaptic “completeness” of our reconstructions, also referred to as the completeness percentage^14^. 17% of synaptic links have an identified neuron on both sides. For other densely reconstructed datasets, this number is: 41.9% for FAFB, 35.3% for HemiBrain, 44.4% for MANC and 40.1% for connections within the neuropils of MaleCNS.

### BANC neuropil mesh generation

We built a “watertight” CNS neuropil surface based on presynaptic sites. We sampled every 500th synapse from the BANC synapse table, and we converted the coordinates to nanometres. To suppress noise, we retained points only if they had ≥10 neighbours within 10 µm (k-NN radius filter). We then fit a 3-D alpha-shape to the filtered cloud (α = 7.5 µm), converted it to a triangular mesh, decimated to ∼25% of faces, Taubin-smoothed (10 iterations), cleaned degenerate/isolated facets, and performed integrity checks. We split the resulting mesh at the neck to yield brain and VNC components. We made each component “integral” (closed) by removing small isolated pieces and isotropically remeshing to fill small gaps; if regions contained multiple pieces, we merged them. We registered neuropil meshes defined in other work into the BANC space (see Neuropils and template alignment). We exported final surfaces as OBJ files, and they are available on our Harvard Dataverse (see Data Availability).

### Synapse prediction evaluation

To determine the false-positive and false-negative rates of the synapse detection, we selected 16 cutouts of size 2 x 2 x 0.7 µm^3^ from across the dataset (in neuropil regions: medulla, lateral accessory lobe, superior posterior slope, gnathal ganglia, fan-shaped body, superior medial protocerebrum, mushroom body medial lobe, mushroom body calyx, anterior ventrolateral protocerebrum, lobula, T1 leg neuromere, wing tectulum, lower tectulum, haltere tectulum, and an anterior and a posterior region of the abdominal neuromere) and compared the predicted synapses with those manually identified by experts (**Extended Data Fig. 1b**). Specifically, the experts identified all synaptic links (pre-post connections) within each cutout without seeing the synapse prediction; two individuals checked each synaptic link. We then compared these manual detections with the predictions and automatically identified true synapses, false positives, and false negatives. Synaptic links for which either the pre or the post was incorrect were considered errors. Finally, the experts examined the synapses in each category and corrected any mistaken or ambiguous calls. We report the number of true, false-positive, and false-negative synapses as well as the F-score, precision, and recall for the individual cutouts and for the dataset as a whole (using the total number of synapses in each category summed across all 16 cutouts). We note that because our synapse detection method relies on identifying postsynaptic profiles, some classes of synaptic connection for which postsynaptic sites are less distinct may be under-detected. We know that our average number of outgoing connections for Kenyon cells (131 links) is far smaller than in FAFB (213 links, cleft score threshold > 50). Another area of under-detection may be axo-axonic connections between sensory neurons. The BANC detection has an autapse rate of 2.1%, a majority of which we expect to be a misassignment of the presynaptic link from a correctly detected postsynaptic link. We recommend users filter out autapses in their analyses.

### Neurotransmitter prediction

We used a recently described approach to predict neurotransmitter type at each automatically predicted synapse^51^. Briefly, we trained a 3D CNN to classify presynapses into one of eight neurotransmitter classes: acetylcholine, dopamine, GABA, glutamate, histamine, octopamine, serotonin, or tyramine. We compiled ground truth data for synaptic transmission from the literature^6,9,66,67,69,70,73,76,124,126,127,136,139,143,146,196–275^, totaling 4416 identified cell types from FAFB/MANC/Hemibrain. Of these, members of 3014 cell types (44153 neurons) could be found in BANC. We removed motor neurons from the ground truth, as they have few presynapses within the CNS. The dataset was split by neuron into training and testing sets, with 80% of the data fortraining and the remaining 20% for testing. We used the following sampling strategy to ensure a balanced dataset across different neuron types. For neurons associated with the most common neurotransmitters (acetylcholine, GABA and glutamate), we randomly sampled a maximum of 10 presynaptic sites from each neuron. For all other neurotransmitters, we included all identified presynaptic sites. This approach ensured that all cell types that had ground-truth were represented in both training and testing sets. The input data for the network consisted of 3D cutouts from the EM volume, each centered on a presynaptic site. These local cutouts had dimensions of 640 x 640 x 630 nm. We used a 3D CNN architecture based on the 18-layer residual network (ResNet-18)^276^. ResNet-18 includes 3D convolutional layers, batch normalization and ReLU activation functions, with the core of the architecture consisting of residual blocks that use skip connections to enable effective training. The model architecture was adapted for our task by modifying the initial convolutional block to accept single-channel grayscale input from EM data. Finally, we replaced the model’s original fully-connected output layer with a linear layer that maps the learned features to our eight specific neurotransmitter classes, followed by a softmax activation to produce the final probability distribution. The network was trained using the Adam optimizer^277^ to minimize the focal loss function^278^. This loss function is a variant of the standard cross-entropy loss, which is effective for datasets with a significant class imbalance as it down-weights the loss assigned to well classified samples, allowing the model to focus on difficult-to-classify samples. To further improve generalization of the model, we applied several data augmentation techniques during training. These included random affine transformations, random noise, and random gamma correction. The probability of applying these augmentations was increased for less frequent neurotransmitter classes to further mitigate the class imbalance. We trained the model for 1,060,000 iterations using a batch size of 16 samples. The final model selected was the one that achieved the highest classification accuracy on the separate testing set. A neuron-level transmitter prediction is obtained by summing the classification probabilities for each predicted class across all presynaptic detections, and selecting the class with the highest total confidence as the most likely neurotransmitter; we assume Dale’s law^279^ holds even though we know that an unknown proportion of neurons in the CNS co-transmit with multiple fast-acting transmitters^51,210,212,280^. Though marginally improved, as in^51^, we expect a large proportion of our serotonin predictions in particular to be incorrect, as the network seems to guess serotonin for peptidergic neurons that lack clear signs for another classification. A fully cited compilation of ground truth labels per cell type can be found here: https://github.com/funkelab/drosophila_neurotransmitters/tree/main, collated by A.S.B., D-Y.A. and J.F.

### Neuropils and template alignment

To transform the BANC data into a standard template space for analysis and inter-dataset comparisons, we computationally generated a ‘neuropil stain’ based on the synapse prediction^195^. To do this, we downsampled and Gaussian blurred (σ = ∼900 nm) the predicted synapse locations to produce a synapse density map at the approximate resolution of light microscopy data used in the Drosophila standard templates. We then registered the synapse density map of the EM dataset to the JRC 2018 Female brain and JRC 2018 Female VNC templates^281^ separately using elastix (https://elastix.lumc.nl/). Leveraging this alignment, neuropils and neurons were transformed between different connectome datasets for visualization and quantitative comparison in the same coordinate system. Meshes for individual neuropils in the central brain^39^ and VNC^40^ were based on previous work. We generated a left-right registration for BANC based on a thinplate-spline warping registration built from matched points on identified pairs of ∼30 DNs, available through the `bancr` R package (`bancr::jrc2 0l8f_to_banc_tpsreg`, `bancr::jrcvnc2 0l8f_to_banc_tpsreg`).

### Proofreading

We proofread neurons to correct automated cell segmentation errors as we described previously^15^. Members of our respective laboratories, dedicated proofreading teams at Princeton, SixEleven (Davao City, Philippines), and Aelysia (Bristol, United Kingdom), as well as a community of citizen scientists collaboratively undertook this effort. We used a multi-pronged strategy. To capture neurons with cell bodies in the CNS, we proofread segments associated with automatically-detected nuclei, which were then extended to reconstruct their full morphology and remove false mergers. To include sensory neurons, whose cell bodies typically reside outside the CNS, we seeded every neuron profile in planes that cut a cross-section through a nerve (1 plane per nerve, except in cases where 1 plane could not capture the full cross-section of the nerve; 47 seed planes total) and then reconstructed starting from those seeds. To capture all neurons in the neck connective, we seeded two planes that were cross-sections through the neck connective (y = 92500 and y = 121000). These transverse planes were positioned posterior to the central brain and anterior to the VNC. Additionally, we proofread orphan segments containing >100 presynaptic links in decreasing order of synapse count for the central brain and VNC. We considered a neuron ‘backbone proofread’ when its primary neurites (if not sensory), or major microtubule-rich processes had undergone a thorough review^36^. This indicated that we expected the overall morphology of the cell to be correct and that, while minor branches or a small number of synapses might still require adjustment, we did not anticipate future proofreading to radically alter the neuron’s core shape or identity. We proofread 114,610 neurons to ‘backbone proofread’. In addition, we defined a further category, “roughly proofread”, to encompass neurons that were identifiable (comprising a large region of arbor), but may have some omission. Frequently, this was due to known dataset artefacts, i.e. regions of the dataset where misalignment or data loss has made reconstruction more challenging. We provide these locations in **Supplementary Data 8**. **Fig. 1c** includes roughly proofread neurons with nuclei in its count and roughly proofread sensory neurons, so as not to potentially count the same cell multiple times (i.e. a roughly proofread orphan axon and its roughly proofread dendrite). In total, 155 people served as proofreaders for the project (defined as individuals who made ≥100 edits).

### Color MIPs

We generated color-depth maximum intensity projections (colorMIPs) of all proofread neurons using the BANC python package (https://pypi.org/project/banc/). We registered neuronal reconstructions to JRC2018_Unisex_20x_HR (1210×566 px) and/or JRC2018_VNC_Unisex_40x_DS (573×1119 px), for compatibility with NeuronBridge^282^.

### Cell-type matching and annotation

#### Overview

Previous studies have invested substantial effort in cell typing both the brain^5,6,14,17,46^ and VNC^7,8,10^, employing a combination of manual annotation and computational methods. Our approach leverages morphology and connectivity matching to cell type the ∼160,000 neurons in the BANC dataset by associating them with published reconstructions and their cell types, namely FlyWire-FAFB v783^15,42^ and MANC v1.2.1^11^. We assigned cell type labels by automatically identifying potential matches between BANC neurons and earlier datasets, based on neuron morphology and position (using NBLAST, **Fig. 1c-e**, **Extended Data Fig. 1e**) as well as connectivity (using NTAC in the optic lobe and network alignment followed by expert review in the VNC). Human annotators then manually reviewed cell type matches. NBLAST alone is competent but not sufficient to make matches; our careful manual review of all neurons in the VNC (human assessment of 3D morphology matches and some integration of known connection information) revealed that 55% of the cell types differ from the type of the max-NBLAST match in MANC. Specifically in the VNC, cell type assignments were more challenging because the reference dataset is of a male fly, and we had to account for sexual dimorphisms, which will be described in A.M., C.K.S. et al., in prep. Nevertheless, we have assigned cell type labels to 64% of BANC neurons (98182 neurons, 93% excluding the optic lobes), with an estimated error rate of ∼7% based on sampling 1,000 matched neurons for more careful review, incorporating cosine similarity to compare neurons between datasets. To do so, connectivity vectors were collapsed by cross-matched cell types up/downstream of each query neuron, and then their cosine similarity was calculated between all query neurons with a mid-to-high NBLAST score between BANC and FAFB/MANC. A majority of mismatches were due to incomplete proofreading; as proofreading improves, matches made improves. The mismatched neuron was almost always a similar cell type within the same hemilineage. For the remaining neurons that could not be confidently matched, we have classified them based on gross morphology and identified their closest associated neurons in other datasets with NBLAST. We estimate that ∼10% of these unmatched neurons will prove unmatchable due to reconstruction quality issues or developmental differences in neuron wiring. Notably, we estimate that as many as ∼892 neurons of the VNC may be sexually dimorphic, ∼4% of the female VNC (A.M., C.K.S. et al., in prep. drawing on the 2025 FlyWire VNC matching challenge: https://codex.flywire.ai/app/vnc_matching_challenge) and cannot be matched well to MANC (which is a VNC sample from a male fly). It has been shown that 2.4% of the female central brain is sexually dimorphic^42^. Our annotation work enables direct comparisons with established identified cell types in the field and facilitates integration with existing datasets. Excluding ANs and DNs neurons (which are truncated in FAFB and MANC), we are missing 679 FAFB cell types in BANC (8%) and 307 cell types from MANC in BANC (11%, but note that MANC is male and BANC is female).

#### Process

Using NBLAST^44^, which quantifies pairwise neuronal similarity by considering both the position and morphology of neuronal arbors and calculating similarity scores by comparing matched morphological segments, we automatically identified potential matches. We matched BANC neurons to reconstructions in FlyWire-FAFB v783^15,17^, MANC v1.2.1^10^, FANC v1116^7^ and HemiBrain v.1.2.1^14^. FAFB is a dense reconstruction of the female brain. MANC is a dense reconstruction of the male ventral nerve cord. FANC is a female ventral nerve cord dataset, with part of the abdominal ganglion truncated, which has been sparsely reconstructed. HemiBrain is a dense reconstruction of ∼½ of the female central brain, including a high-quality reconstruction of the mushroom body and central complex. Recently updated^42^ cell type names from FAFB were preferred for brain matches, from MANC for VNC matches, FANC for known female-specific^21^ ANs and motor neurons^8^ (where the annotation is better than in MANC) and HemiBrain for the central complex^6^ (where the annotation is better than in FAFB). Following automated NBLAST scoring, we manually reviewed candidate matches. For sensory neurons, ANs and DNs and intrinsic neurons of the brain, this manual review involved co-visualizing the meshes of matched neurons in three orthogonal 2D projections and evaluating the correspondence. For ANs and DNs, we followed up this 2D comparison with co-visualization and manual evaluation in 3D using neuroglancer. For intrinsic neurons of the VNC, we also used connectivity to automatically determine their similarity to MANC neurons. When the top matched cell type agreed between NBLAST and connectivity, we assigned the neuron to that cell type; when these potential matches were in conflict, we co-visualized the BANC and MANC neurons in 3D in neuroglancer and manually reviewed them to determine the correct cell type. High NBLAST scores (e.g., above 0.3) generally indicated a strong likelihood of a correct match. Iterative proofreading and matching increased the population of identified cells as sometimes, low NBLAST scores indicated issues with neuron reconstruction, which suggested additional proofreading was necessary.

For many afferent (i.e. sensory) and efferent (i.e. effector including both motor and endocrine) neurons, in addition to matching to FAFB and MANC, we used comparisons to the literature and the domain expertise of our authors to determine their cell types and functions. In particular, we identified leg and wing motor neurons by their morphology and connectivity, as previously described^8^. The key identifying features we used were the exit nerve of the axon, the relative trajectory of the primary neurite, the relative position of the soma, and unique features of the dendritic morphology. Front, middle, and hind limb neuropils differ in terms of specific morphology yet the identifying motor neuron features largely retain their relationships, allowing us to identify homologous motor neurons in each neuropil^9^. We confirmed morphological identification by comparing these motor neurons on the basis of the sources of common synaptic input^8^. We identified endocrine neurons of the brain based on morphology and the cosine similarity of their connectivity with each other and with the FAFB endocrine neurons. We used morphological comparisons to the literature to identify the motor neurons of the antennae, eyes, neck, crop, haltere, pharynx, proboscis, pharynx, salivary glands and uterus; octopaminergic effector neurons involved in ovulation; endocrine neurons of the VNC; and chemosensory, tactile and proprioceptive sensory neurons from the head, eyes, antennae, proboscis, legs, abdomen, wings and halteres^283^. In some cases, we used data from the larval fly (putative nociceptive, putative oxygenation and aorta sensory neurons^10,61,63,64,284–287)^ to annotate suspected homologous neurons. Adult nociceptors will be reported^60^. We subjected chordotonal, campaniform and hair plate neurons of the VNC, including those of Wheeler’s organ, the prothoracic organ and the metathoracic organ, to additional careful review and re-annotation^7,79,288–290^. We could not identify chordotonal and bristle neurons of the haltere based on the literature; these are a minority input from the halteres.

#### Neurons of the neck connective

We reviewed all profiles in the two seed planes through the neck connective. We successfully proofread 98.3% of the neuronal profiles to ‘backbone proofread’ status, for a total of 3695 proofread neurons. We then matched these neurons to cell types in FAFB and MANC, as described above, except for the 8 female-specific DN cell types and 31 female-specific AN cell types identified in ^21^, which we instead matched to FAFB and FANC. We identified 1839 ANs, of which we matched 1722 (corresponding to 548 cell types), and 1313 DNs, of which we matched 1294 (corresponding to 478 cell types). In addition, we identified 13 sensory DNs (afferent axons that enter through a brain nerve and project through the neck connective to the VNC, discussed in more detail here^21^) corresponding to 6 cell types, 512 sensory ANs (afferent axons that enter a VNC nerve and project through the neck connective to the brain) corresponding to 42 cell types and 5 efferent ANs (ANs that also project out of other nerves) corresponding to 3 cell types, including EAXXX079, which may be the leucokinin ANs in^291^. We identified all 16 of the female-specific DNs (corresponding to 8 cell types) and 36 of the expected 70 female-specific ANs (corresponding to 16 of the 31 cell types). For ANs, sensory ANs and efferent ANs, we use the MANC cell type name except for the female-specific ANs, where we use the FANC name; for DNs and sensory DNs, we use the FAFB name. When this resulted in the same name for different cell types (which became apparent when considering the full neuron rather than just the brain or VNC half), we appended an underscore and a letter to the FAFB/MANC name. We also identified and proofread 49 efferent neurons of the neck that leave through the cervical nerve. These are neck motor neurons, and we named them as in^58^. Note that because they do not traverse the entire extent of the neck connective, they are not included in our count of 3723 “backbone proofread” neurons of the neck connective. We do not use sensory or efferent ANs and DNs in our analysis of ANs and DNs. In our review of the neck connective, we identified 31 ANs and DNs that appeared to have developed abnormally or were stochastic in whether they had an ascending/descending arbor. For example, DNge079 on the left side (in MANC named DNxl080) has a mis-targeted dendrite located in the VNC, rather than the central brain. However, we note that both the left and right IN08B003 neurons are ANs in this dataset, but are intrinsic neurons of the VNC in MANC and in FANC. We determined that the cell type DNg28 leaves the brain through the maxillary-labial nerve and after it re-enters through the same nerve, its processes remain outside of the glial sheath surrounding the CNS as it then traverses the neck to envelop the outside of the VNC and target neurohemal release sites. Therefore, we re-classified it from a DN to solely an efferent cell type. As in FAFB, we could not find DNg25, and DNd01 was not a DN but rather a central brain intrinsic neuron^21^. Important prior work bridged a proportion of ANs and DNs between FAFB and MANC using available experimental data^21^, which was a valuable resource of our matching efforts.

#### Annotation taxonomy

We annotated neurons hierarchically by flow relative to the CNS (afferent - inputs the CNS, intrinsic - contained within the CNS, efferent - leaves the CNS), super class (eg. sensory, motor, visceral/circulatory, ascending, descending), cell class (eg. chordotonal organ neuron, leg motor neuron, kenyon cell), cell sub class (eg. wing steering motor neuron, front leg hair plate neuron, PPL1 dopaminergic neuron), individual cell type, and with associated metadata (region, side, nerve, body part sensory, body part associated with effector neurons, peripheral target type, cell function, cell function detailed, hemilineage, neurotransmitter verified, neuropeptide verified, FAFB v783 match ID, MANC v1.2.1 match ID and other names). The full list of terms used in each category are listed in **Supplementary Data 1**. This framework enabled both broad and fine-grained categorization, such as distinguishing different and specific classes of sensory neurons. We imported annotations from cell type matching to existing *Drosophila* connectomes^10,15,17^ as well as those that proofreaders and the community contributed through a custom Slackbot (https://github.com/jasper-tms/the-BANC-fly-connectome/blob/main/slackbots/annotation_bot.py) directly to CAVE, facilitating real-time tagging and collaborative refinement. We updated annotations as proofreading progressed, and they are publicly available through FlyWire Codex and on CAVE (cell_info and codex_annotations tables). We have used cell_info for community annotations, and codex_annotations for centralised annotations federated by the core BANC team. We used SeaTable internally to collect, draft and stage our annotations before uploading to FlyWire Codex and CAVE.

Elsewhere in the literature, ‘effector’ is often understood as denoting a muscle or gland. Here we use ‘effector neurons’ to describe motor neurons, endocrine cells, and efferent neurons targeting the viscera. All these cells effectuate change outside the CNS, and thus they are the relevant output units of the CNS.

#### Influence

The influence score^92^ quantifies the influence of a neuron or group of neurons, called the seed, on each of the other neurons in the network. It is a measure of steady-state signal activation, resulting from continuous stimulation of seed neurons. We compute steady-state activation assuming a linear dynamical model,

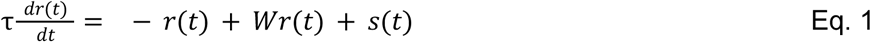

where *r* is the vector of neural activity, *W* is the connectivity matrix, τ is the network time constant, and *s* is the simulated neural stimulation. For each seed, all elements in s corresponding to the seeded neurons are set to one, while the remaining elements are fixed at zero.

The weight of each connection is taken as the number of synapses in that connection, normalized by the total count of input synapses onto the postsynaptic cell in question. That is, if *c_ij_* is the synapse count from presynaptic neuron *j* onto postsynaptic neuron *i*, then the total input count for neuron *i* is

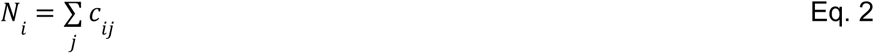

and the connectivity weights were set to

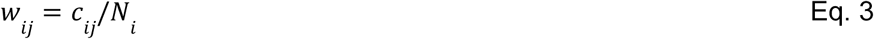

This type of normalization follows previous work and has been shown to qualitatively capture experimental observations^46,292^. Connections with fewer than 5 synapses were filtered out in the analysis. All connectivity weights are treated as nonnegative values. To ensure stable neural dynamics, we re-scaled *W* such that its largest real eigenvalue is 0.99.

We compute the steady-state solution for the assumed network dynamics by

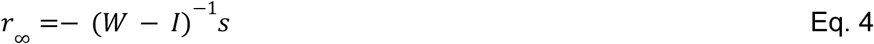

separately for each seed vector *s*. As *W* is a highly sparse matrix, we could compute this solution efficiently using the sparse matrix parallel computing libraries PETSc and SLEPc (https://petsc.org/release/ and https://slepc.upv.es/).

If the seed is one cell, and we are interested in a single target cell, we simply take the steady-state activity of the target *r* in response to the seed. We define *r_ij_* as the steady-state response of target cell *j,* given stimulation of seed cell *i*. To better understand the steady-state solution, note that using the Von Neumann series, it can be rewritten as an infinite sum of powers of the connectivity matrix *W* multiplied by the seed vector s,

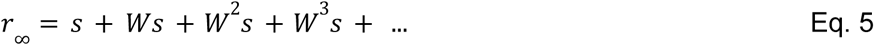

Here, the first term is the immediate impact of the seed(s), and the remaining terms quantify its impact across immediate synaptic connections - or one ‘hop’ - across two hops, and so on. This shows that we are summing over seed propagations across different ‘hops’ until all direct and indirect pathways between seed and target neurons have been covered.

Often, we are interested in pools of related target cells (e.g., a pool of related motor neurons). Thus, for a target pool *T* that contains the indices of the |T| = *N* target neurons, we take the average steady-state response of each cell in the target pool

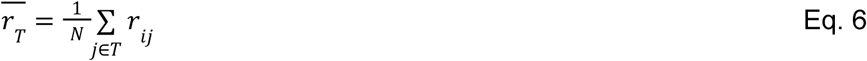

Similarly, we are often interested in a pool *S* of related seed cells, where *S* contains the seed cells’ indices. Here, we could simulate activity in all seeds individually, and average the results. In this case, for a seed pool of size |S| = *M,* the average response is

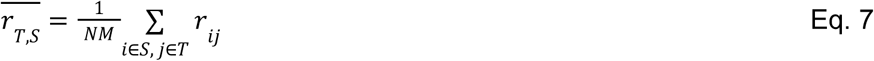

Alternatively, because the steady-state solution *r*_∞_ is linear in the seed vector, it is sometimes more convenient to just simulate activity in all seed cells simultaneously. In this case, if *r_j_* is the response of the *j*th target cell to the simultaneous activity of all seed cells, we take

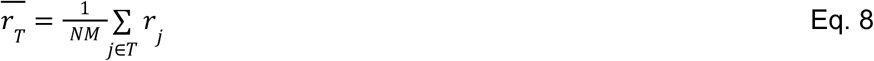

In this type of simulated network, *r̅* will generally decay exponentially as the distance increases between the seed and the target (in network space). To correct for this, we take the logarithm of *r̅*; note that *log(r̅*) should be generally proportional to the number of hops separating the seed and the target. And because *log*(*r̅*) is generally negative, we add a constant *c* that brings the values of *log*(*r̅*) into the nonnegative range, for ease of display.

The resulting value is called the “adjusted influence”:

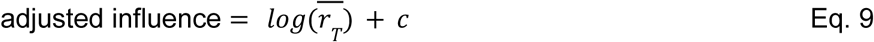

In a few cases (only as noted in the corresponding figure legends), we calculate adjusted influence, corrected for seed size:

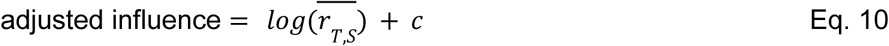

This is because, in these cases, we are interested in quantifying the total influence of a sensory cell sub class onto effector neurons. In all other cases, we are interested in quantifying the influence between the average source and target neuron of the given classes, and so use the first formula.

We used *c* = 24, because this ensured that all adjusted influence values were non-negative (given that −24 was approximately the minimum value of *log*(*r̅*) we observed). Across the entire CNS, a small and discrete group of cells had *log*(*r̅*)<<-24 for any seed, as these cells were not well-connected to the graph; we set these adjusted influence values to 0.

We confirmed that adjusted influence is approximately inversely proportional to the number of synaptic ‘hops’ separating the seed cells and target cells, as expected, and this was true for two different published metrics of hop length (**Fig. 2b**. and **Extended Data Fig. 4a**; see below for details of these previous metrics). Thus, adjusted influence is essentially a computationally efficient and deterministic method of estimating a quantity that is proportional to the effective number of hops separating the seed and the target. Because the number of hops is an unsigned quantity^17,22^, it is reasonable that adjusted influence is also unsigned. Moreover, we do not actually know the signs of all connections in the connectome, particularly concerning glutamatergic, dopaminergic, serotonerergic, octopaminergic, and tyraminergic synapses. Notably, even inhibitory cells can drive behavior, e.g. via disinhibition; the only output of the vertebrate cerebellar cortex consists of GABAergic Purkinje cells, which trigger motor output when they pause firing, thereby disinhibiting their downstream targets^293^.

As compared to previous metrics of hop number, adjusted influence has several advantages. First, we have an explicit expression for the steady-state solution, making the computation more efficient relative to comparable activity propagation approaches^15,22,46^. Second, the steady-state solution is linear in the seed vector, such that it can in principle be summed across different seeds.

Rather than taking the steady-state activity as the basis for this influence metric, we also considered using the initial slope of the neural activity. However, the initial slope turned out to be directly proportional to the chosen seed vector, which made it unsuitable as a measure to quantify network-wide influences. We furthermore considered projections of the above dynamics into the top 1000 eigencircuits, similar to previous work^87^, but we found this truncation to be unsuitable for our purposes to well-approximate the full network dynamics.

The reader should note that, as with the synaptic count between two neurons, the adjusted influence between two neurons is best understood relative to adjusted influences calculated between comparable sources and targets. There is no value over which an important interaction is assured.

We computed influence scores reported in this paper using Python 3.13.2, and we executed all computations using a MacBook Pro running macOS Monterey version 12.6.9. The code used to compute the influence scores is available as a separate Python package^92^ (see ‘Code availability’ section).

#### Alternative metrics of polysynaptic connectivity

For comparison with our influence scores, we used two complementary probabilistic graph traversal algorithms to model information flow through the CNS. First, we applied the signal cascade approach^22^, in which activity propagates from a set of seed neurons to downstream targets based on synapse counts, treated as proxies for synaptic strength. A key feature of this model is that neurons are activated only once and then enter a deactivated state, enabling assessment of potential temporal sequences of activation.

Second, we used an information flow model^15,46^, in which neurons are probabilistically recruited based on the fraction of synapses received from already recruited neurons. This model allows ongoing activation from previously active neurons and assigns each neuron a rank that reflects its integration point in the circuit. While these ranks do not correspond to true physiological latency, this approach enables systematic inference of information flow directionality and network layering across the CNS.

#### Betweenness centrality

To quantify how strongly each neuron acts as a bottleneck in the BANC, we computed each neuron’s betweenness centrality, a standard network metric that quantifies how often a given node lies on the shortest paths between pairs of other nodes (excluding the endpoints)^294–297^. Neurons with high betweenness centrality lie on many routes between other neurons and are positioned at putative bottlenecks in the network.

We calculated two different versions of betweenness centrality: all-to-all betweenness and sensory-to-effector betweenness. To calculate all-to-all betweenness, we computed the unnormalized, unweighted, directed betweenness centrality for all the neurons in the network using the Brandes algorithm^295^ as implemented in the Python interface of igraph^298^ (**Fig. 3a**, **Extended Data Fig. 7a**).

We represented the network as a directed graph *G* = *(V, E),* where *V* is the set of neurons and *E* is the set of directed edges representing connections between neurons. We included a directed edge only if the connection had at least 5 synapses. The all-to-all betweenness of a neuron *v* is defined as follows^295^:

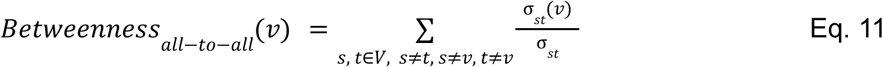

where *σ_st_* is the number of shortest paths from s to *t* and *σ_st_*(*v*) is the number of shortest paths from *s* to *t* that include *v.* If there are multiple shortest paths, 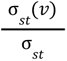 indicates the fraction of such shortest paths that include neuron v. We then sum this fraction over all pairs of neurons *s*, *t* ∈ *V* with *s ≠ t,* and the neuron *v* is neither the source nor the target (*s ≠ v*, *t ≠ v*). Since our graph is directed, paths from s to *t* must follow edge directions.

To quantify how strongly each neuron acts as a bottleneck in the sensory to effector information flow, we calculated the source-target betweenness centrality^296^. Specifically, we used the same unnormalized, unweighted, directed betweenness centrality but restricted the source–target pairs to sensory neurons (“sensory ascending”, “sensory descending”, and “sensory”) as sources *S* and effector neurons (“motor”, “visceral circulatory”, “ascending visceral circulatory”) as targets *T* when accumulating betweenness scores (**Fig. 3a**).

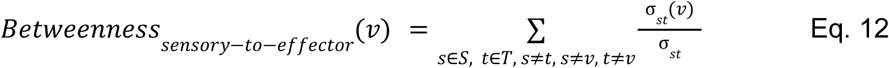

#### Spectral clustering

We adapted a spectral clustering algorithm^299^ to partition the CNS into modules of highly interconnected cells. For this analysis, we focused on intrinsic neurons of the central brain and VNC, ANs, DNs, visual projection neurons, and visual centrifugal neurons. (We chose to exclude optic lobe intrinsic neurons because they are so numerous that they end up dominating the analysis.) Starting with these 42,639 cells, we iteratively pruned cells that did not have at least one input and output partner among the remaining cells (e.g. because all their input comes from sensory neurons, or all their output goes to motor neurons, etc.). This left 41,951 cells as the input to this analysis.

To apply spectral clustering, we first specified our population of *N* cells of interest and a desired number of clusters *k.* We then constructed a weighted, undirected graph whose nodes corresponded to these *N* cells and whose edge weights were derived from the connectome. More formally, edge {*i*, *j*} was assigned weight

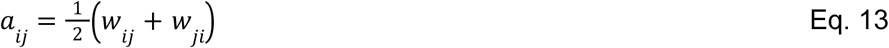

where *w_ij_* the normalized synaptic input from presynaptic cell *j* to postsynaptic cell l, as defined above. We then computed the first *k* (*k* = chosen cluster number) eigenvectors of the graph Laplacian, which resulted in a *k × N* matrix of unit-norm eigenvectors *X*. Each node then received a k-dimensional feature vector that was determined by its loadings onto the eigenvectors, yielding an *N × k* feature matrix *Y* with entries

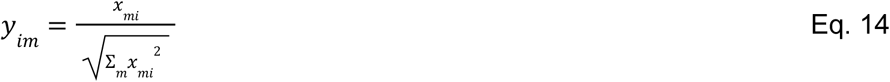

Finally, we applied k-means clustering to these feature vectors to assign each node to a cluster. We decided to use 13 clusters because this produced a coarse-grained division at the approximate level of resolution we found relevant to our analysis, and also because the resulting cluster divisions largely corresponded to salient boundaries in the UMAP space of CNS neurons.

### Data analysis

Visual projection neuron functions were used to account for different visual information streams as ‘sensors’. This is an incomplete survey of visual functions bounded by the literature^129,143,145–158^. We used for visual_chromatic - aMe12, MeTu3b, MeTu3c, MTe50; visual leg feedback - LT52; visual horizontal wide field motion - dCH, FD1, FD3, H1, LPT04_HST, LPT21, LPT22, LPT23, LPT26, LPT42_Nod4, Nod1, Nod2, Nod3, vCH; visual large_objects and visual thin vertical bar - LC15; visual loom - LC16, LC4, LPLC1; visual object and visual loom - LC12, LC17; visual polarized light - MeMe_e10, MeTu2a, MeTu2b, MeTu3a; visual small_object - LC10a, LC10b, LC10c, LC10d, LC11, LC13, LC18, LC21; visual small_object,visual_loom - LC26, lC6, LC9, LPLC2; visual thin vertical bar - LC25, MeTu1; visual vertical wide field motion - LPT27, LPT28, LPT30, LPT31, LPT45_dCal1, LPT47 vCal2, LPT48 vCal3, LPT49, LPT50, Nod5, V1, vCal1, VST1, VST2. Sensory neuron cell functions were determined by a literature search and search of extant connectome metadata, for information on their peripheral sensory organs/structures. Through this manuscript, we clustered heatmaps using hierarchical clustering based on Ward’s distance using functions from base R. We applied dynamic tree cut^300^ (implemented as dynamicTreeCut::cutreeDynamic, using deepSplit = 4) clustering to UMAPs to delineate effector and AN/DN clusters, other than in **Fig. 6 and Extended Data Fig. 10**, in which spectral clustering was used, see above. We conducted data analysis in R using the *uwot*^301^, *tidyverse*^302^ and *ggplot2*^303^ packages. We made the Kernel density estimates for **Fig. 6a** using MASS::kde2d, n=100, cubes with densities above the first percentile colored^304^. We calculated cosine similarity using the lsa R package^305^, and we applied it to direct connectivity between BANC neurons to build the space used in **Fig. 3**. To perform the Kolmogorov-Smirnov test in **Fig. 6e,f**, we used the *kstest2* function in MATLAB 2024a (Mathworks) and to perform two-way ANOVA tests and Kruskal-Wallis tests, we used their respective statistical functions using the *car* and *stats* packages in R. P-values for non-parametric Wilcoxon rank-sum and signed-rank tests were obtained using the *wilcox.test* function in R. Significance asterisks: **** p<0.0001, *** p<0.001, ** p<0.01, * p<0.05 and ns p>0.05. We used LLM assistance to review and recommend code as well as to draft code documentation, all of which we consciously evaluated for accuracy and which was in compliance with the Harvard University Generative AI guidelines (https://www.huit.harvard.edu/ai/guidelines). The Harvard AI Sandbox (https://www.huit.harvard.edu/ai-sandbox) provides a secure environment in which to use LLMs, and all queries are recorded. The majority of our codebase was not assisted by LLMs.

## Data availability

Data is freely accessible through multiple platforms. A general overview of the resource and links to these tools are available at the BANC portal (https://banc.community). FlyWire Codex^306^ (https://codex.flywire.ai/banc) provides an interactive web interface for exploring the BANC connectome, enabling users to search for neurons, visualize morphology, traverse synaptic pathways and download metadata such as cell-type annotations, neurotransmitter predictions and connectivity matrices. Volumetric EM data, including 3D neuron meshes and annotations, can be viewed at https://ng.banc.community/view or accessed programmatically via CAVE^37^. We snapshotted CAVE materialization version 626 (July 21,2025) for this manuscript. Static data and code dumps are also available for download from a DOI-minting repository, the Harvard Dataverse (DOI: https://doi.org/10.7910/DVN/8TFGGB). Direct downloads include: the synaptic connectivity edgelist, NBLAST results of BANC neurons against Hemibrain, FAFB, FANC and MANC as well as BANC all-by-all; neuronal L2 skeletons (made using: https://github.com/CAVEconnectome/pcg_skel); neuronal colorMIPs; influence scores; our aligned BANC metadata; template registrations; behavioral data for the BANC fly; and snapshots of our code. Aligned image data is available on BossDB (DOI: https://doi.org/10.60533/boss-2025-941r, or URL: https://bossdb.org/project/bates_phelps_kim_yang2025). Schematics are available here as vector graphics: https://github.com/wilson-lab/schematics?tab=readme-ov-file. Some code and analyses in this manuscript (i.e. for **Fig. 1**) used a more recent materialization, version 745, which the reader can find on the public google bucket: `gs:/sjcabs_2025_data/banc/`. On publication, Codex will reflect the latest version of the data.

## Code availability

All code developed for this project is open-source and publicly available. Our connectome data is most accessible through FlyWire codex, where it is browsable and from where up-to-date direct downloads can be obtained, as the project progresses (https://codex.flywire.ai/banc). A comprehensive collection of community tools and software packages for working with the BANC dataset can be found at the project hub (https://banc.community) and the FlyWire Apps portal (https://flywire.ai/apps). The specific code used to perform the analyses and generate the figures for this manuscript is shared in a dedicated GitHub repository: https://github.com/htem/BANC-project/. Code for computing influence scores is available at: https://doi.org/10.5281/zenodo.17693838 for python and at https://github.com/natverse/influencer for an R implementation. Code for neurotransmitter predictions is available at: https://github.com/htem/synister_banc. We have also made available python code for BANC (https://pypi.org/project/banc/), and an R package, `bancr` (https://github.com/flyconnectome/bancr), for querying BANC data, compatible with the natverse^83^. A static snapshot of the code and analysis tools are also available on our Harvard Dataverse Dataset (DOI: https://doi.org/10.7910/DVN/8TFGGB).

## Extended Data Figures

**Extended Data Fig. 1:**
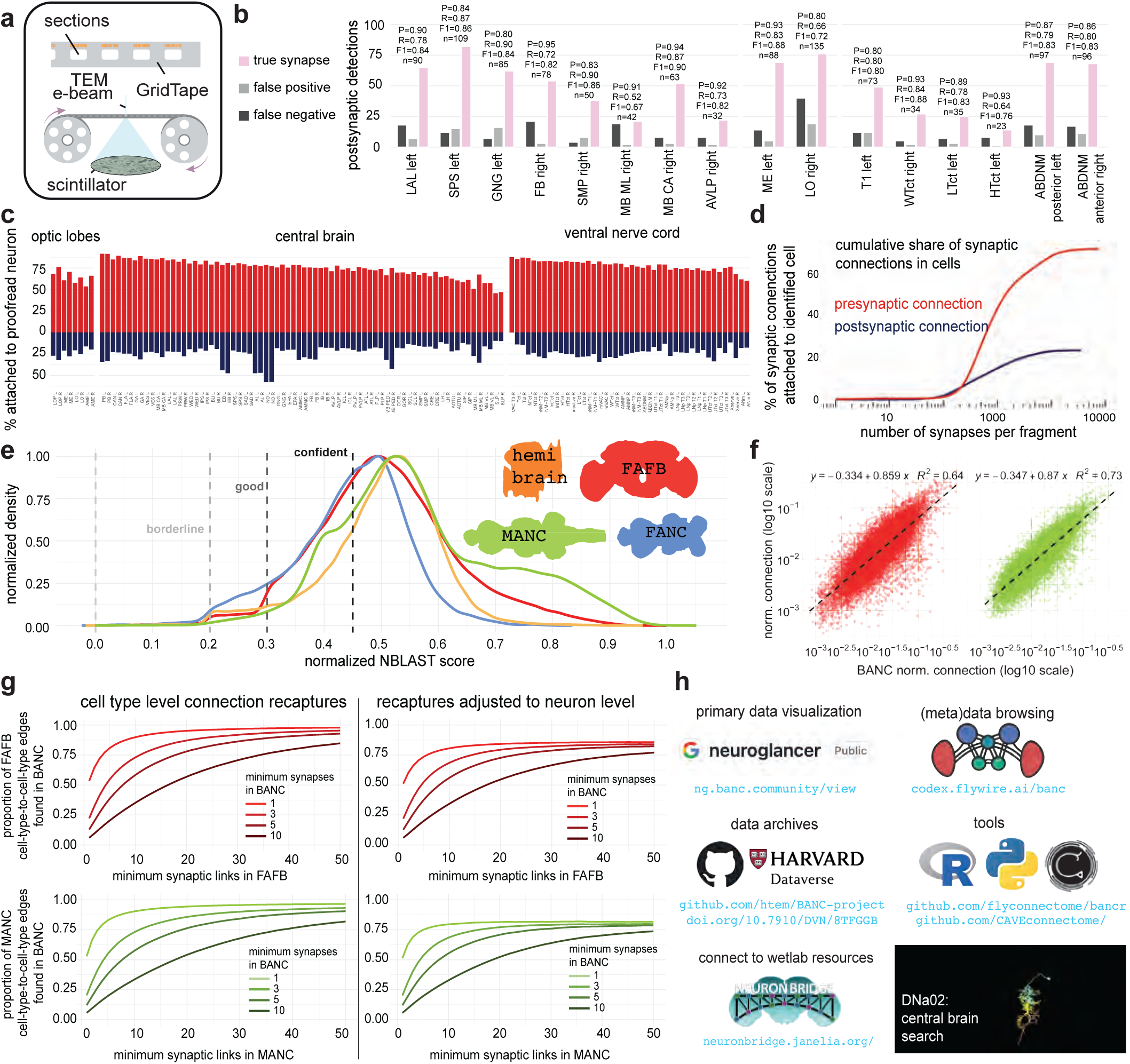
Central nervous system connectome generation, quality and neuron identification. a. Overview of the GridTape^7^ imaging approach. Serial EM sections are automatically collected onto a tape-based substrate and subsequently imaged in a transmission electron microscope (TEM) with a custom reel-to-reel imaging stage. b. We evaluated postsynaptic predictions in 16 neuropil cutouts (P, precision; R, recall; F1, F1-score). c. Attachment rates for presynaptic (red) and postsynaptic (navy) links to an ‘identified’ neuron (backbone proofread and roughly proofread neurons/fragments, known non-neuron and known glial synaptic detections removed) across neuropils (174598 cells). We used the BANC synapse version: synapses_v2 and BANC segmentation and metadata version: v821. d.The cumulative share of pre- and postsynaptic links in identified neuron versus orphan fragments (not part of an identified neuron). Plot is by fragment size as inferred by number of links on fragment (version 626). e. Density of the normalized NBLAST scores^44^ of ‘backbone proofread’ neurons^15^ in the BANC against all neurons in other connectomic datasets (different colors). We consider normalized NBLAST scores > 0.3 as high and suggest score bins to help guide data users (dashed lines). Normalized NBLAST scores are “raw” NBLAST scores divided by the self-match score. All density curves are normalized to their own peak. FAFB is a dense female brain reconstruction. MANC is a dense male VNC reconstruction. FANC is a sparse female VNC reconstruction. HemiBrain is a dense reconstruction of ∼½ of the central brain. f. Scatter plots show the correlation between matched pairs of connected cell types in the BANC versus FAFB^15^ and MANC^11^ (the most complete extant connectomes). Each point is a cell-type-to-cell-type normalized connection (synaptic connections from source-to-target / total number of postsynaptic links on the target cell type). FAFB-BANC: 34174 matched cell type connections, MANC-BANC: 29350 matched cell type connections. g. Rates of connection recapture between the FAFB (upper) and MANC (lower) datasets and BANC. Both cell type level connections (left), and an adjustment for neuron-level connections (right) are shown. The y-axis shows the proportion of edges recaptured in BANC. Cell-type pairs are treated as presence/absence (no size normalization). Only cell types present bilaterally in both datasets and their common edges are evaluated. On the right, each e.g. FAFB cell-type pair contributes a density-based weight to approximate neuron-level recovery. For each source–target cell-type pair we: 1. compute its connection density in FAFB (neuron–neuron links divided by the product of the numbers of neurons in the two types). 2. compute the same density in BANC. 3. take the BANC-to-FAFB density ratio, capped at 1, and use that as the pair’s contribution. The curve shows the average of these contributions across all FAFB pairs that meet the x-axis threshold - i.e., the mean neuron-level recovery of known FAFB/MANC connections in BANC. h. Users can browse BANC data via Codex (codex.flywire.ai/banc), and they can download data for programmatic analysis (via Codex^15^, CAVE^37^, and the Harvard Dataverse at https://doi.org/10.7910/DVN/8TFGGB). Color-depth MIPs^307^ (maximum intensity projection images where color encodes depth) in JRC2018U space^281^ for BANC dataset neurons (version 626) are available from the Dataverse archive. These can be used to search for genetic driver lines enabling functional investigation into BANC neurons, for example using NeuronBridge^282^. Example search image shown for a specific cell type (DNa02).

**Extended Data Fig. 2:**
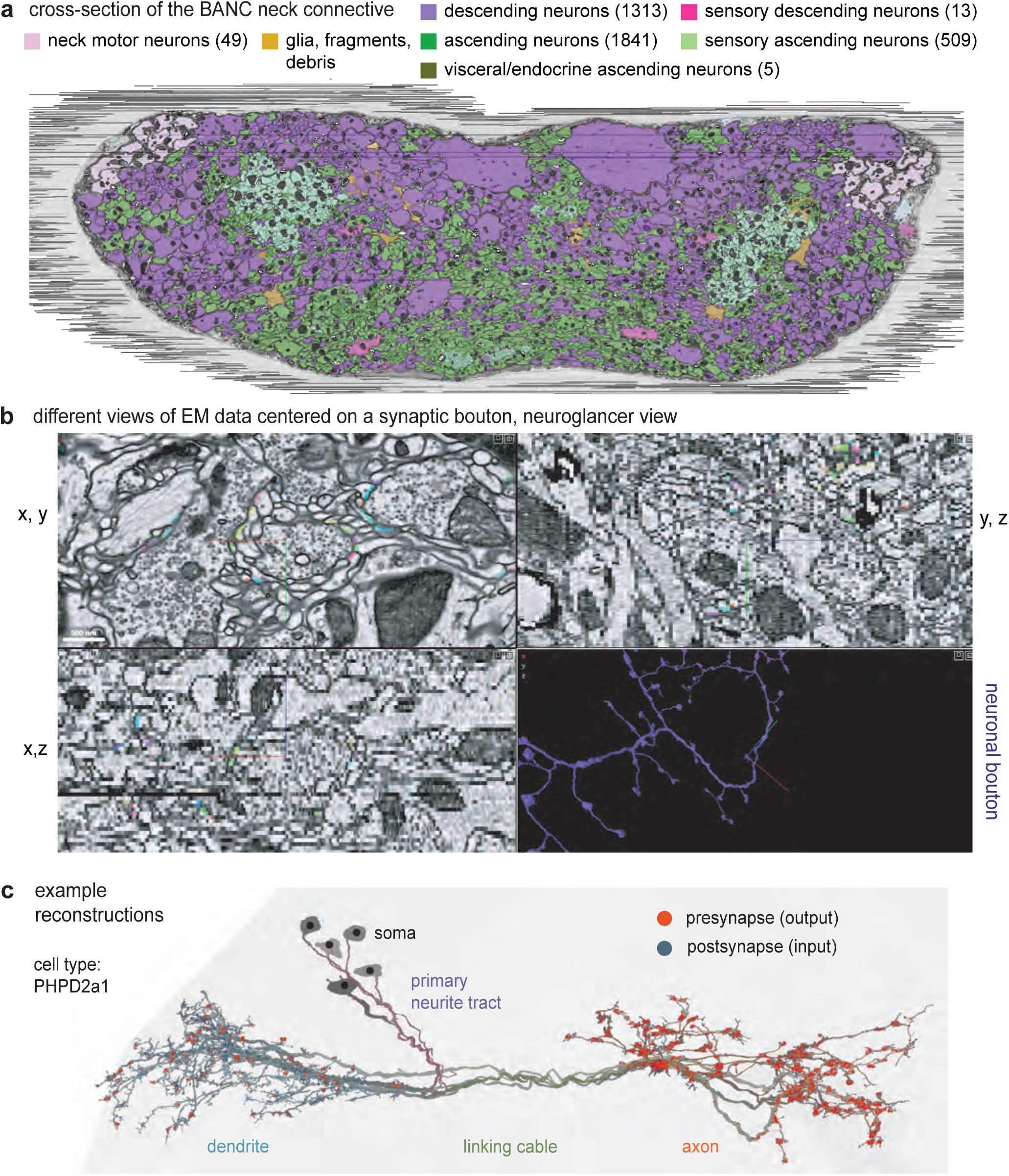
Neck connective neurons and example reconstructions. a. Cross-section of the neck connective from BANC, neuronal segmentations shown in color by super class labels. b. Different views of cross-sections around a synaptic bouton in neuroglancer, shown at 4 x 4 x 45 nm, in xyz. Note that the public instance downscales data to 8 x 8 x 45 nm in xyz. Lower right, lines point to the bouton location in the neuronal arbor. c. Example reconstructions of identified LHPD2a1^308^ neurons (grey meshes), with skeletonizations (lines), root points (black dot), synapse detections (blue = input, red = output) and compartment determinations^309^ shown.

**Extended Data Fig. 3:**
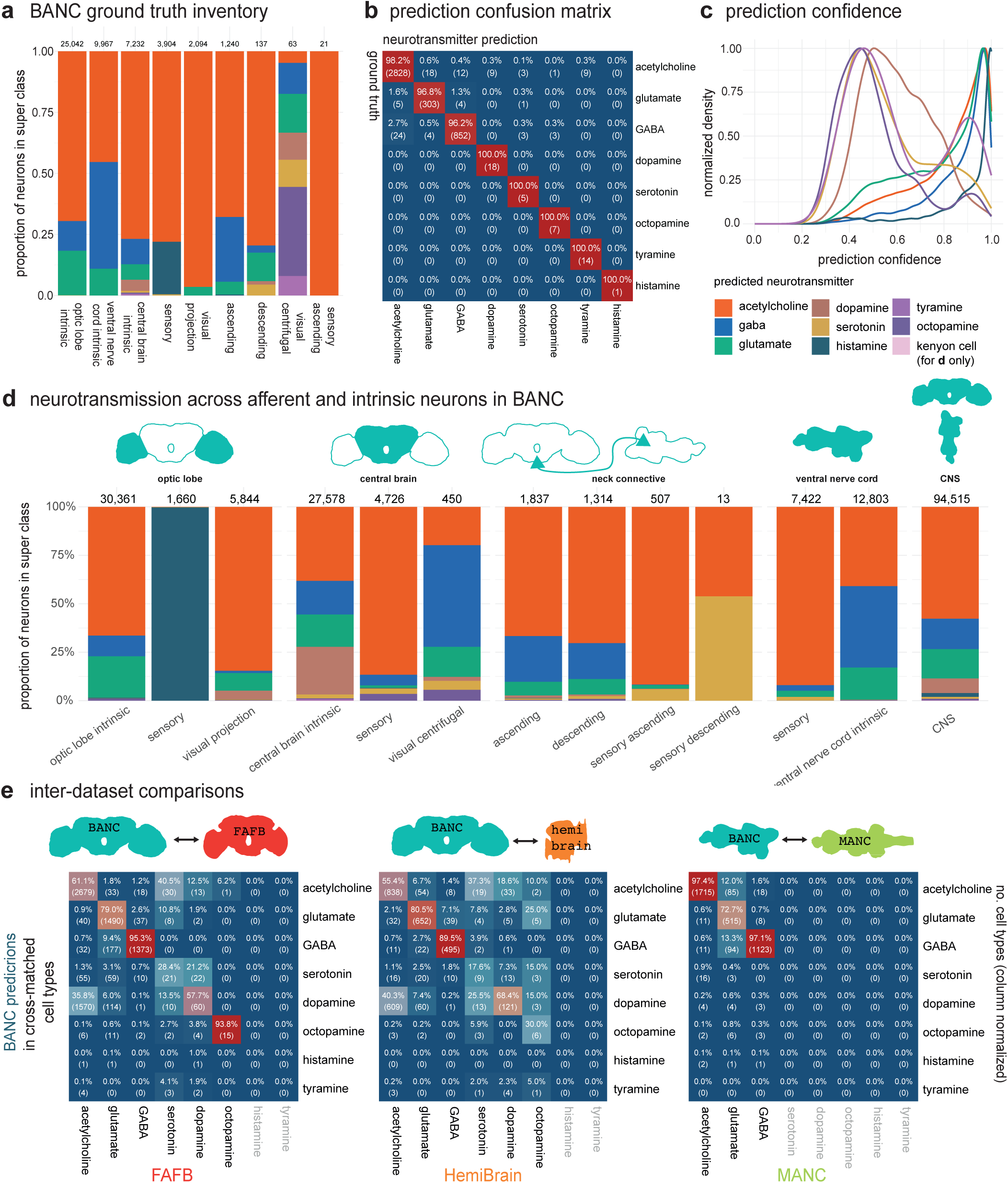
Expanded neurotransmitter predictions across the CNS. a. Stacked proportions of verified neurotransmitters per neuron super class (one verified transmitter per cell type from literature). Effector neurons are removed, as their presynaptic links are mostly outside of the dataset. b. Confusion matrix of neurotransmitter prediction evaluated at the level of whole neurons on the held-out test set. Whole neuron prediction is based on the summed classification probabilities across all presynaptic links, selecting the most confident class. The ground-truth included 20572 neurons (from 2900 cell types, see Methods), of which 16448 were used for training and 4124 for testing. c. Normalized density of neuron-level prediction scores (the winning proportion of classified presynaptic links that determined the neuron-level prediction) across proofread, non-effector neurons. d. Stacked proportions of predicted neurotransmitters within each super class plus a single CNS summary bar for all BANC proofread neurons with at least 50 presynaptic links. Effector neurons are removed, as their presynaptic links are mostly outside of the dataset. Kenyon cells are mostly mispredicted as dopaminergic, as in FAFB and HemiBrain^51^ and so are shown in a separate color for clarity. e. Cell-type modal neurotransmitter comparison between BANC (rows) and each external dataset (columns): FAFB^51^, Hemibrain^51^, MANC^11^. Only cell types present in both datasets are included. Greyed out transmitters were not predicted in the given dataset.

**Extended Data Fig. 4:**
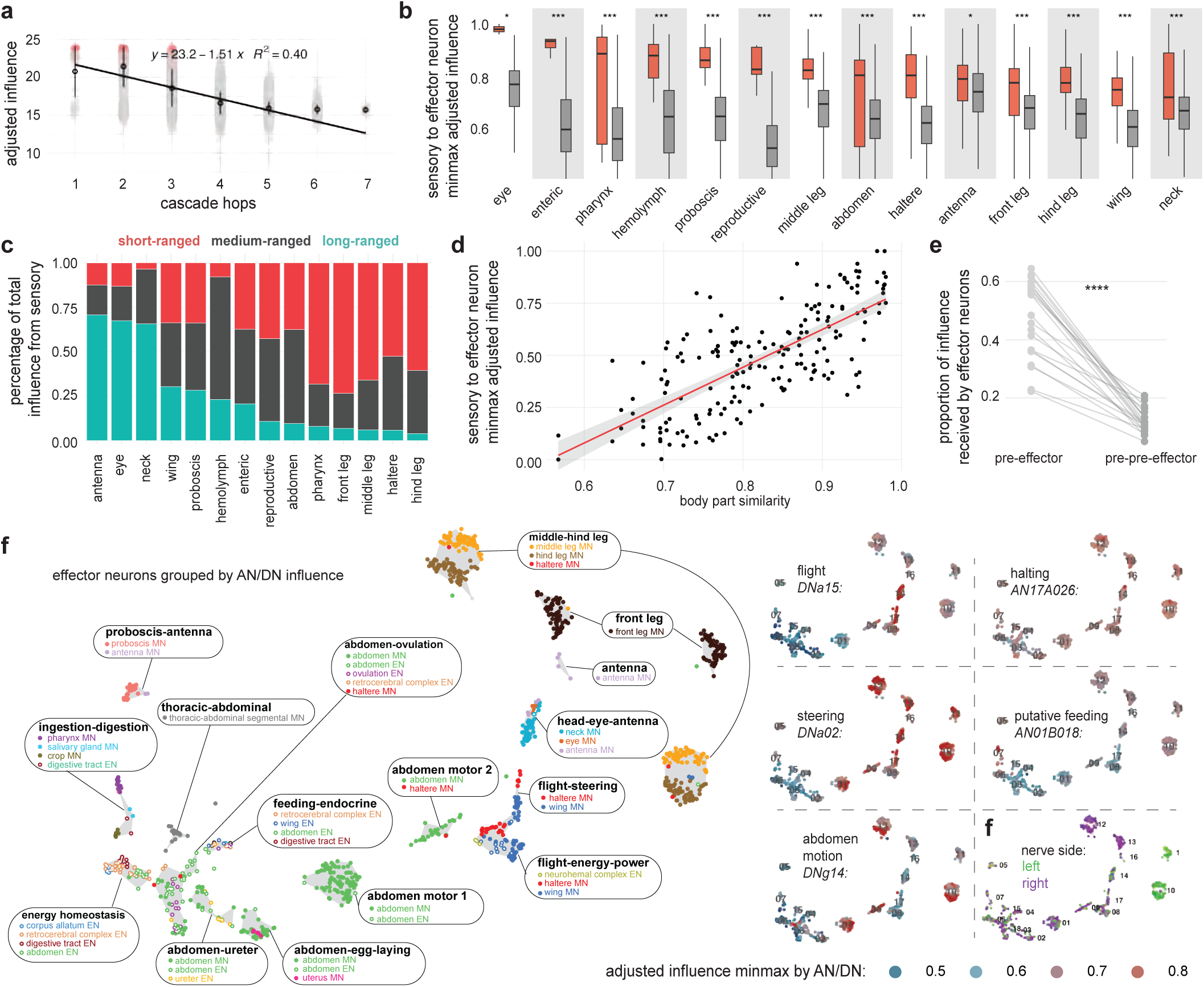
Extended analysis of influence. a. Fig. 2b shows that the adjusted influence is proportional to ‘layers’ of a published graph traversal model^46^ applied to the FAFB dataset^17^. Here we show that the adjusted influence is also proportional to the output of a different published layering algorithm^22^. As in Fig. 2b, we used olfactory seeds annotated in the FAFB dataset. b. Boxplots for each body part (n=14 body parts) comparing adjusted influence (Eq. 10) values from sensory sub classes to effector neurons, grouped by whether the sensory and effector body parts are the same (matched) or different (unmatched). We normalized by target, as in Fig. 2e. Wilcoxon rank-sum tests (one-sided: matched > unmatched, Holm-corrected (α = 0.05) across all comparisons) show that matched sensory-effector pairs have significantly higher influence across all body parts (* p < 0.05, *** p < 0.001). c. Division of sensory influence onto effector neurons (grouped by body part). Sensory influence is categorized as local (same body part), mid-range (sensory and effector neurons are both in the brain or both in the VNC but have different body parts), or long-range (sensory neuron is in the brain and effector neuron is in the VNC or vice versa). d. Scatterplot shows normalized, adjusted influence values from Fig. 2e, i.e. between effector body part pairs (n=196 pairs from 14 body parts) versus the mean cosine similarity between effector body parts (i.e. calculated between the columns in Fig. 2e). Same body part pairs have cosine similarity = 1 by definition and are omitted. Pearson correlation shows a significant positive relationship between body part similarity and influence (r = 0.729, R^2^ = 0.531, p = 1.97×10^-31^). e. Proportion of influence effector neurons of a body part receive from pre- and pre-pre-effector neurons of their matched body part. “Pre” indicates neurons one direct synaptic connection upstream of an effector neuron (synaptic connections > 5 synaptic links); “pre-pre” indicates neurons two direct synaptic connections upstream of an effector neuron. Proportion of influence becomes more distributed from pre-effector to pre-pre effector neurons (paired Wilcoxon signed-rank test, p = 0.954×10^-6^). f. Left side: map of effector neurons based on the AN/DN influence that they receive. Note, for example, that DNs and ANs co-regulate pharynx motor neurons, salivary gland motor neurons, crop motor neurons, and endocrine neurons of the digestive tract; these effector neurons are labeled “ingestion-digestion”. Similarly, DNs and ANs coordinate wing power motor neurons, haltere motor neurons and endocrine neurons of the neurohemal complex; these effector neurons are labeled “flight-energy-power”. As these examples illustrate, DNs and ANs often co-regulate effector neurons in different body parts. Right side: the same effector neuron maps, color-coded by adjusted influence from example ANs and DNs. Minmax normalization rescales influence values to 0-1, across all AN/DNs. This illustrates the influence distribution rather than magnitude. Green and purple: the same effector cell map, color-coded by the side of the CNS on which their efferent axon exits.

**Extended Data Fig. 5:**
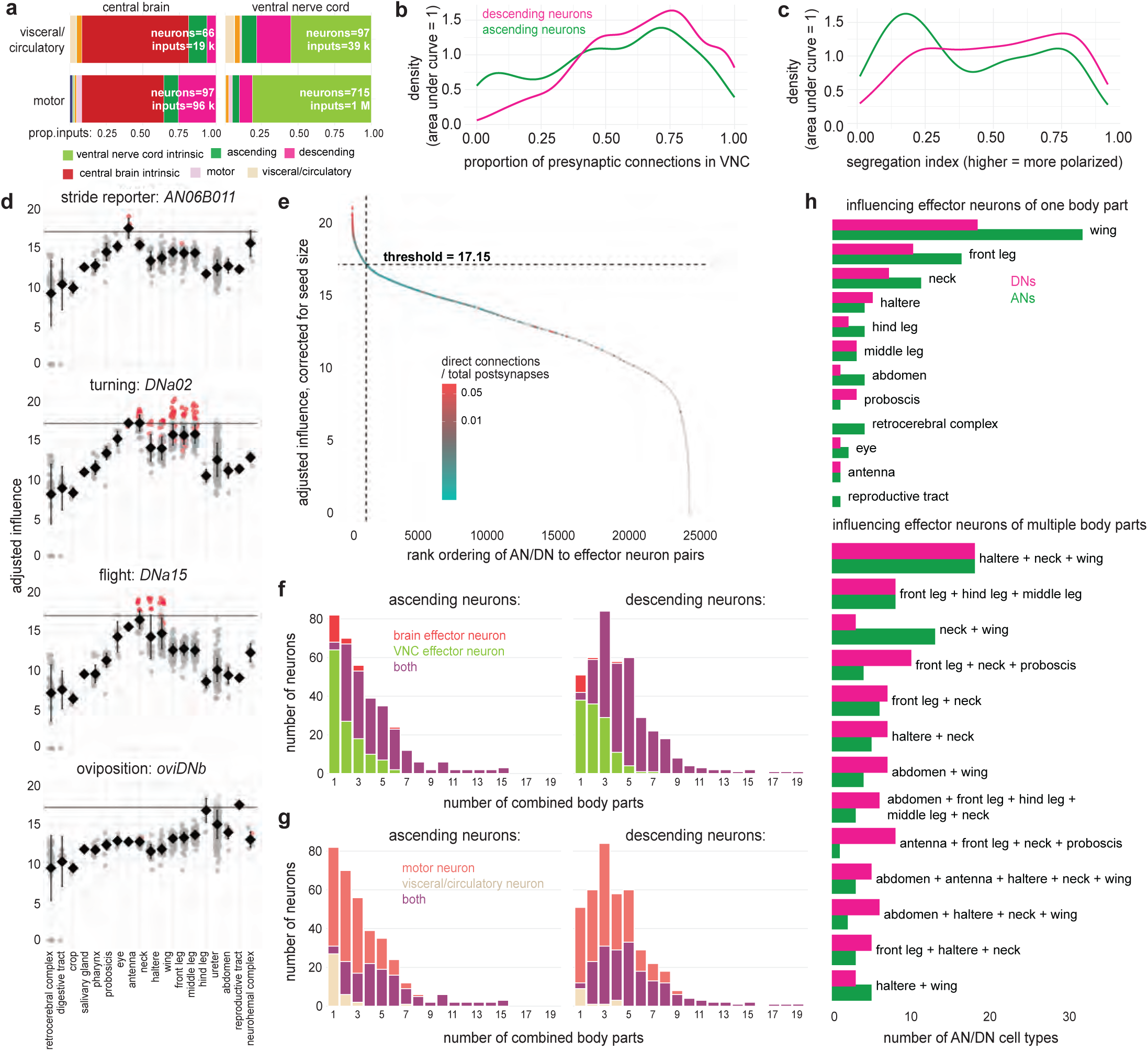
Individual DNs and ANs often influence effectors in multiple body parts. a. The proportion of input synaptic links to effector neurons by the super class identity of the presynaptic neurons. *Top*, visceral/circulatory neurons, *bottom*, motor neurons. *Left*, effector neurons in the central brain, *right*, effector neurons in the VNC. b. Distribution of presynaptic links in the VNC versus the brain, for all DNs (1313 cells) and ANs (1841 cells) in the BANC dataset. c. Distribution of segregation index^309^ values for these same DNs and ANs. Segregation index is a measure of polarization that quantifies the entropy of pre- and postsynaptic connections between the axonal and dendritic compartments of a neuron. A segregation index closer to 1 indicates a more polarized neuron. d. Here we chose three DNs and one AN that have clear behavioral effects, and we examined their adjusted influence on effector neurons in different body parts. Within each subplot, each point is an effector neuron, with direct connections in red. The horizontal line marks a value of 17.15, which we take as a conservative cutoff for “high influence” (see note below). All four cells have some effector neuron influence above this cutoff. For each cell, the above-cutoff effector neuron influences are compatible with the cell’s function. e. The adjusted influence cutoff as the “elbow” in the cumulative distribution of AN/DN-to-effector neuron adjusted influences (Eq. 10) involving DNs and ANs with known behavioral functions; DNs and ANs used to identify this elbow were DNa02^110^, DNa01^110^, DNp01^121^, DNp02^122^, MDN (DNp50)^102^, DNp42^109^, DNg97^103^, DNg100^103^, DNg12^107^, DNg62^104^, DNp07^106^, DNp10^106^, DNg14^101^, DNa15^120^, DNb01^120^, DNp37^139^, oviDNb^76^, DNp20^113^, DNp22^113^, DNp25^310^, DNp44^310^, DNp2 7^245^, AN17A026^114^ and AN19A018^103^. f. The number of AN and DN cell types that combine different numbers of body parts, after indirect connections are binarized by the cutoff in (e). Gross CNS division for combined effector neurons shown in color (‘both’ can appear when only one body part is targeted because neck motor neurons are found in both the brain and VNC^58^). g. Same as (f), but color indicates whether the combinations incorporate motor neurons, visceral/circulatory neurons or both. h. After discarding connections below the cutoff in (e), we counted the number of AN and DN cell types that influence effector neurons in single body parts (top) or multiple body parts (bottom). The bottom plot shows only the most common 20 combinations of body parts.

**Extended Data Fig. 6:**
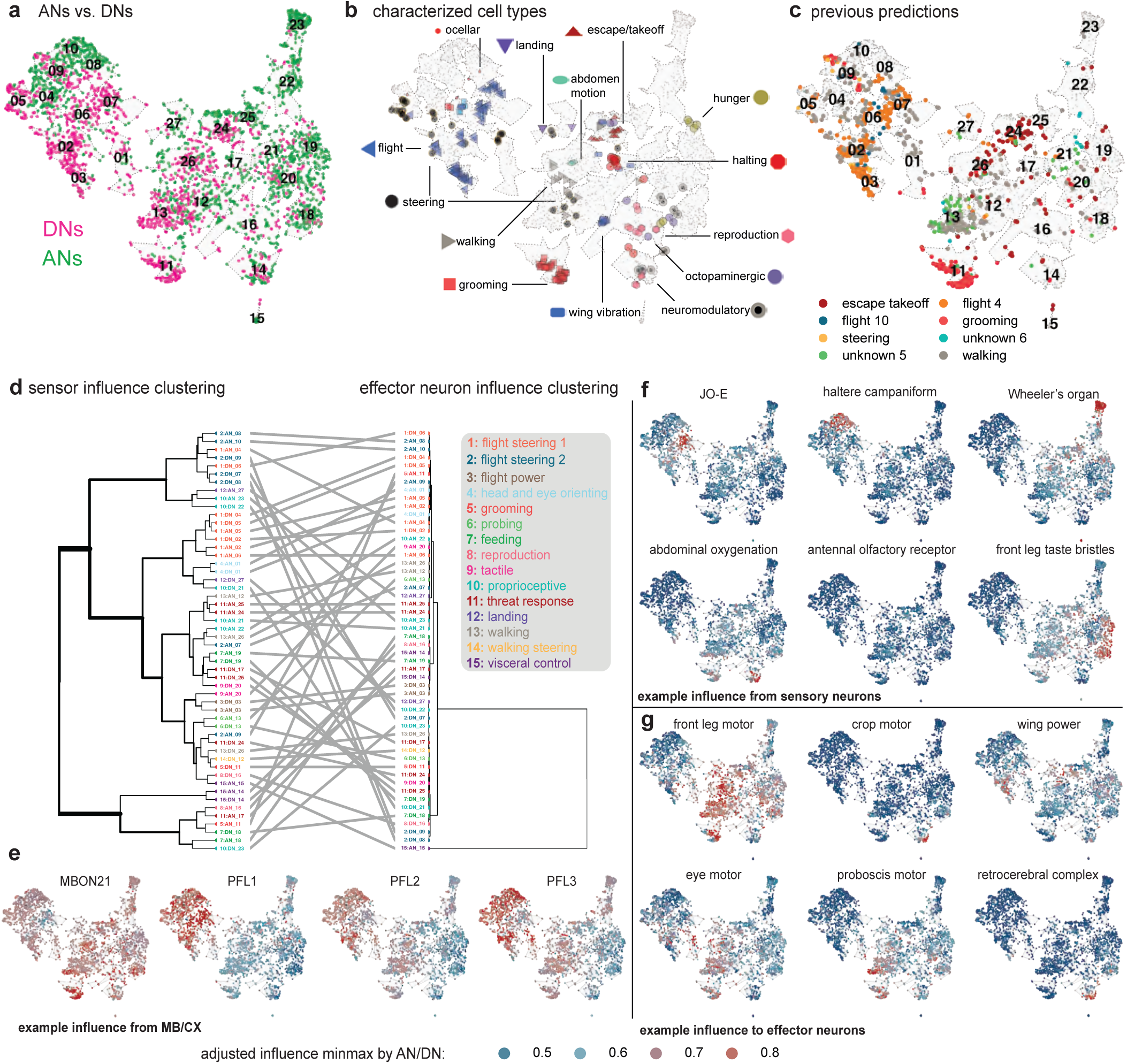
Influence streams to and from AN/DN clusters. a. The UMAP from Fig. 3d (built by AN/DN direct connectivity with other neurons of the CNS); each point is a single AN or DN, colored accordingly. b. Behavioral functions for functionally characterized cell types, mapped onto the same UMAP space. c. The same UMAP with functions assigned by Braun et al. (2024)^101^. This earlier work only used direct FAFB DN-DN connectivity, and as a result, functional information was more limited than it is now. d. Tanglegram showing the relationship between two methods of sorting AN/DN clusters (**a**). The left dendrogram sorts clusters based on the similarity of their adjusted influences from sensory neuron sub classes. The right dendrogram sorts clusters based on the similarity of their adjusted influence to effector neuron sub classes (right). Colors denote superclusters. e. Example normalized, adjusted influence (Eq. 10, source size corrected) from neurons from more ‘cognitive’ brain regions, the mushroom body (MB) and central complex (CX). Minmax normalization rescales influence values to 0-1, across all AN/DNs. This illustrates the influence distribution rather than magnitude. f. Example normalized, adjusted influence (Eq. 10) from sensory neuron sub classes. g. Normalized, adjusted influence (Eq. 10) from ANs/DNs to example effector neuron classes.

**Extended Data Fig. 7:**
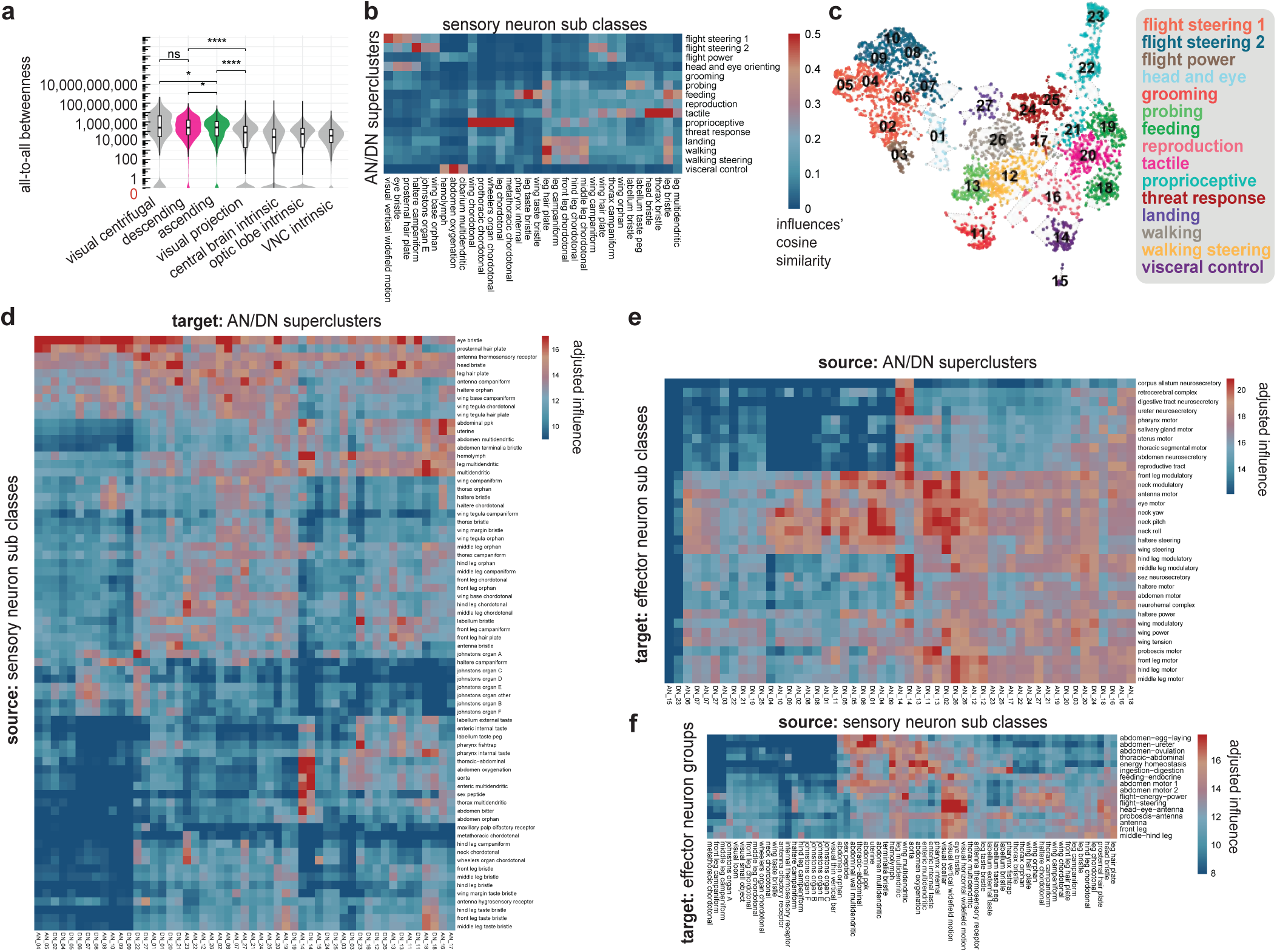
Evidence for AN/DN assignment to behavioral ‘superclusters’. a. All-to-all betweenness by super class in the BANC dataset. For a neuron, its score is the number of shortest paths between designated sources and targets (in this case, all other proofread neurons in BANC v626) that pass through that node. High-betweenness neurons are bridges linking otherwise separate modules. Critically, a node can have low degree but high betweenness if it sits between communities. I.e. this metric captures a neuron’s potential control over global flow, not just its local connectivity. All-to-all betweenness varies significantly with super class (Kruskal-Wallis test, p<2.2×10^-16^). Post-hoc pairwise comparisons using Dunn’s test with Holm correction (α=0.05) revealed that ANs and DNs have significantly higher all-to-all betweenness than other super classes with a lower median (all p < 1.26×10^-71^). Pairwise Wilcoxon rank-sum tests with Holm correction (α=0.05) between ascending and descending neurons and their neighbouring super classes in this plot, showed significant differences between: ascending vs descending (p=0.0305), ascending vs visual centrifugal (p=0.0305), ascending vs visual projection (p=1.78×10^-75^), descending vs visual projection (p=2.34×10^-72^) but not descending versus visual centrifugal (p=0.333). Boxes show a middle line for the median, box ends = 25th and 75th percentiles of non-logged betweenness. b. Cosine similarity between sensory neuron sub classes (columns) and AN/DN superclusters. c. The UMAP from Fig. 3d, with clusters demarcated as numbers. Clusters were determined using a dynamic tree cut algorithm^300^. These clusters were amalgamated into higher level ‘superclusters’ (colors) based on review of their received influences from sensory sources (c), and their influence upon effector neuron sub classes (d), as well as by reviewing where AN/DN cell types of known function fall (**Extended Data Fig. 6**b). d. Adjusted influence (Eq. 10, source size corrected to control for large variation in the size of source classes) from all sensory neuron sub classes onto AN/DN clusters from (b). e. Adjusted influence (Eq. 10) from AN/DN superclusters onto effector neuron sub classes. f. Adjusted influence (Eq. 10) from sensory neuron sub classes onto effector neuron groups from **Extended Data Fig. 4**f.

**Extended Data Fig. 8:**
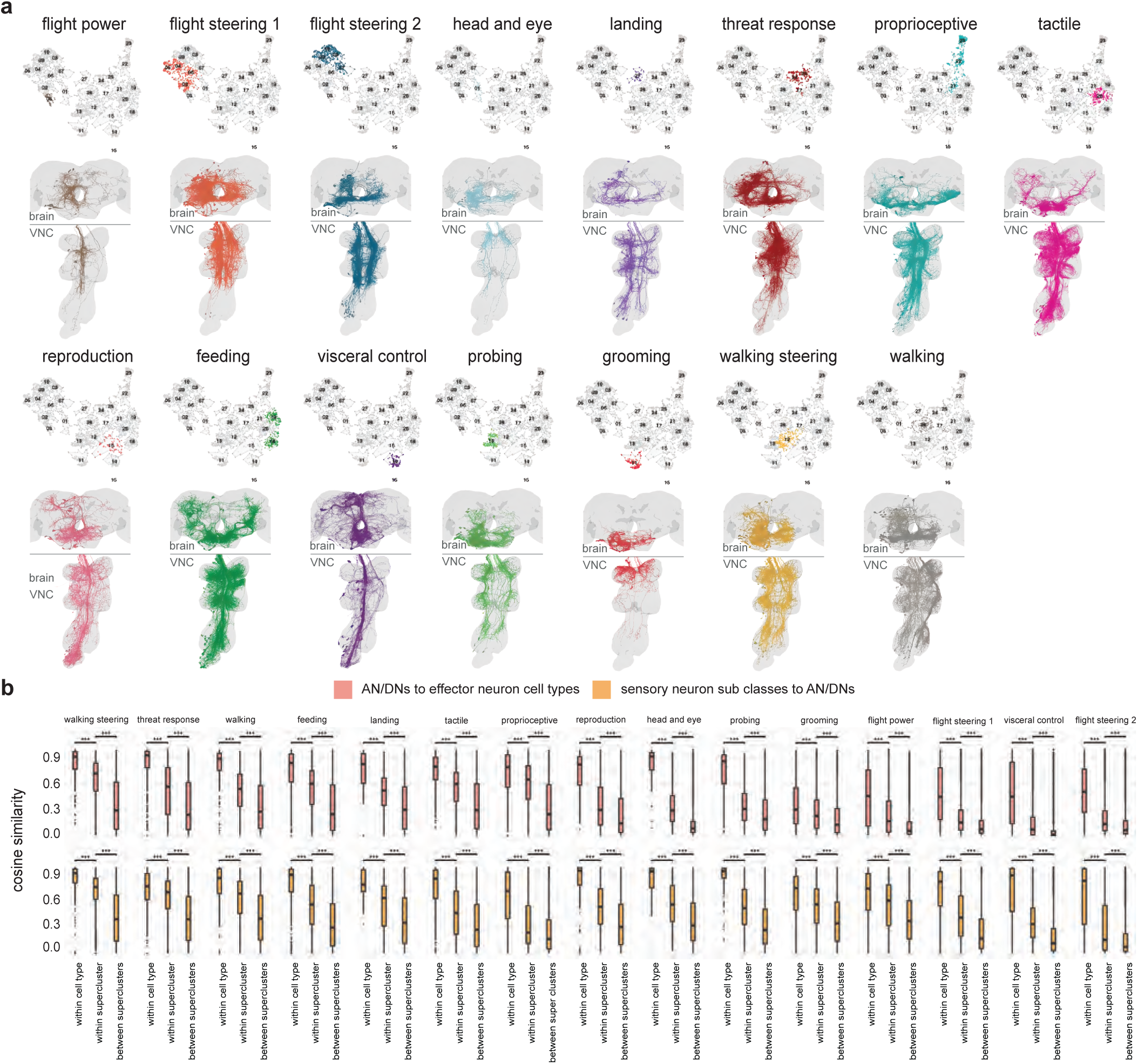
AN/DN morphologies by supercluster. a. Each subpanel shows all right (soma) side neurons from one AN/DN supercluster in the UMAP embedding (Fig. 3d). Neuroglancer links for [flight power](https://ng.banc.community/2025a/flight-power), [flight steering 1](https://ng.banc.community/2025a/flight-steering-1), [flight steering 2](https://ng.banc.community/2025a/flight-steering-2), [head and eye orienting](https://ng.banc.community/2025a/head-and-eye-orienting), [landing](https://ng.banc.community/2025a/landing), [threat response](https://ng.banc.community/2025a/threat-response), [proprioceptive](https://ng.banc.community/2025a/proprioceptive-perception), [tactile](https://ng.banc.community/2025a/tactile-perception), [reproduction](https://ng.banc.community/2025a/mating-and-reproduction), [feeding](https://ng.banc.community/2025a/feeding), [visceral control](https://ng.banc.community/2025a/visceral-state), [probing](https://ng.banc.community/2025a/probing), [grooming](https://ng.banc.community/2025a/grooming), [walking steering](https://ng.banc.community/2025a/walking-steering) and [walking](https://ng.banc.community/2025a/walking). b. Cosine similarity in “from sensory” and “to effector” influences for AN/DN superclusters. Similarity distributions are shown for within cell type, within supercluster and between supercluster comparisons; significance tested with pairwise Wilcoxon rank-sum tests (Holm-corrected across all comparisons, α = 0.05), **** p<0.0001.

**Extended Data Fig. 9:**
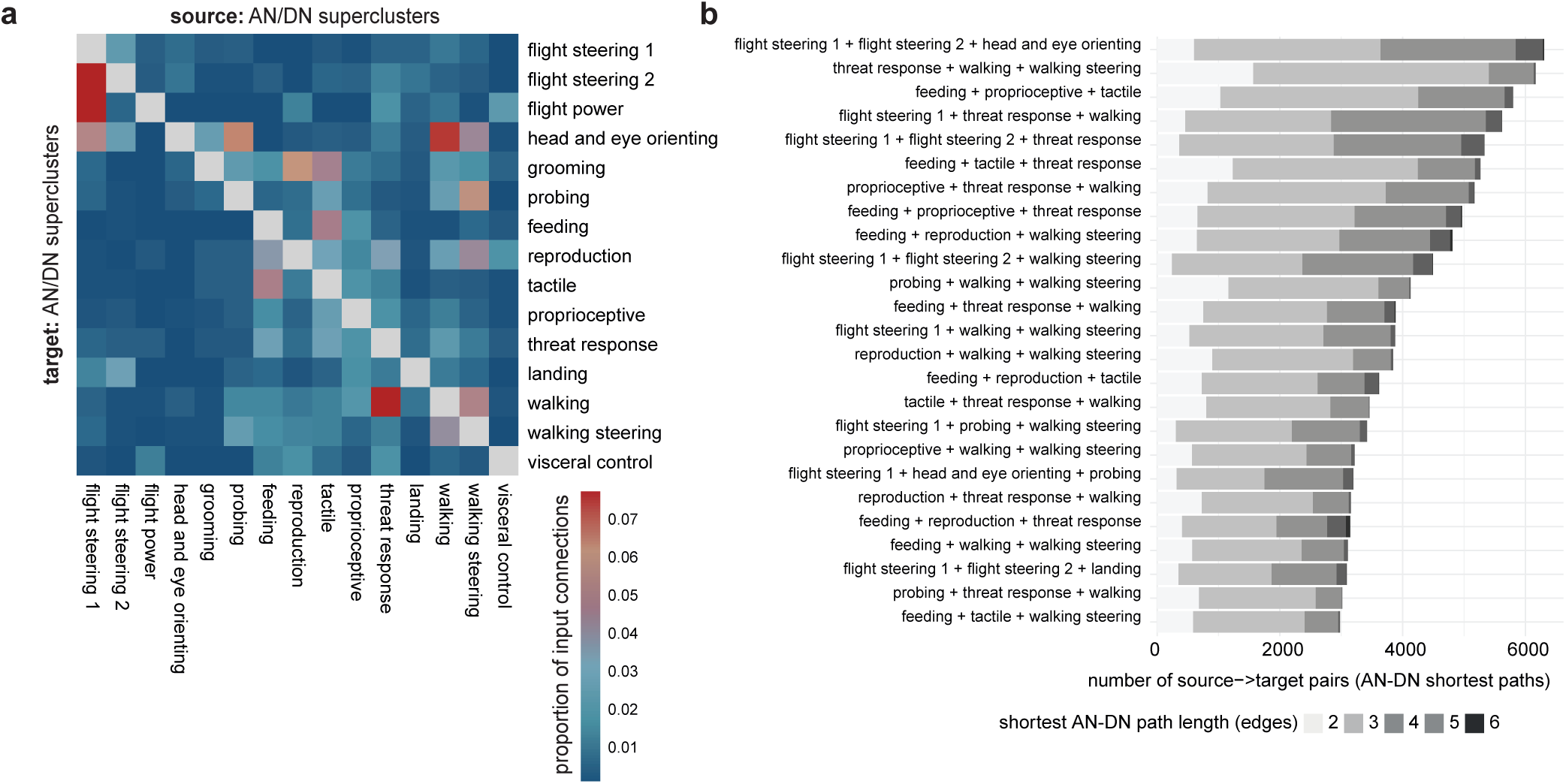
Interactions between AN/DN superclusters. a. Direct connections between AN/DN superclusters, shown as the proportion of each supercluster’s total input budget supplied by the upstream supercluster. b. The number of shortest paths between ANs/DNs in an AN-DN only network (i.e. the BANC edgelist subsetted to only ANs and DNs, threshold ≥10 synaptic links) that transit between 3+ superclusters. Bars show the number of source→target cell-type pairs (AN/DN) whose shortest path traverses ≥3 superclusters. Each bar corresponds to a unique supercluster combination; paths are deduplicated and simple (no node repeats). The 25 most common 3+ supercluster transits are shown. These short-path connections linking 3+ superclusters are akin to motifs reported in the larval *Drosophila* CNS^22^. Note that there are many two-hop chains passing between three different superclusters. The vignette in Figure 5c is one such example, from the most common motif here.

**Extended Data Fig. 10:**
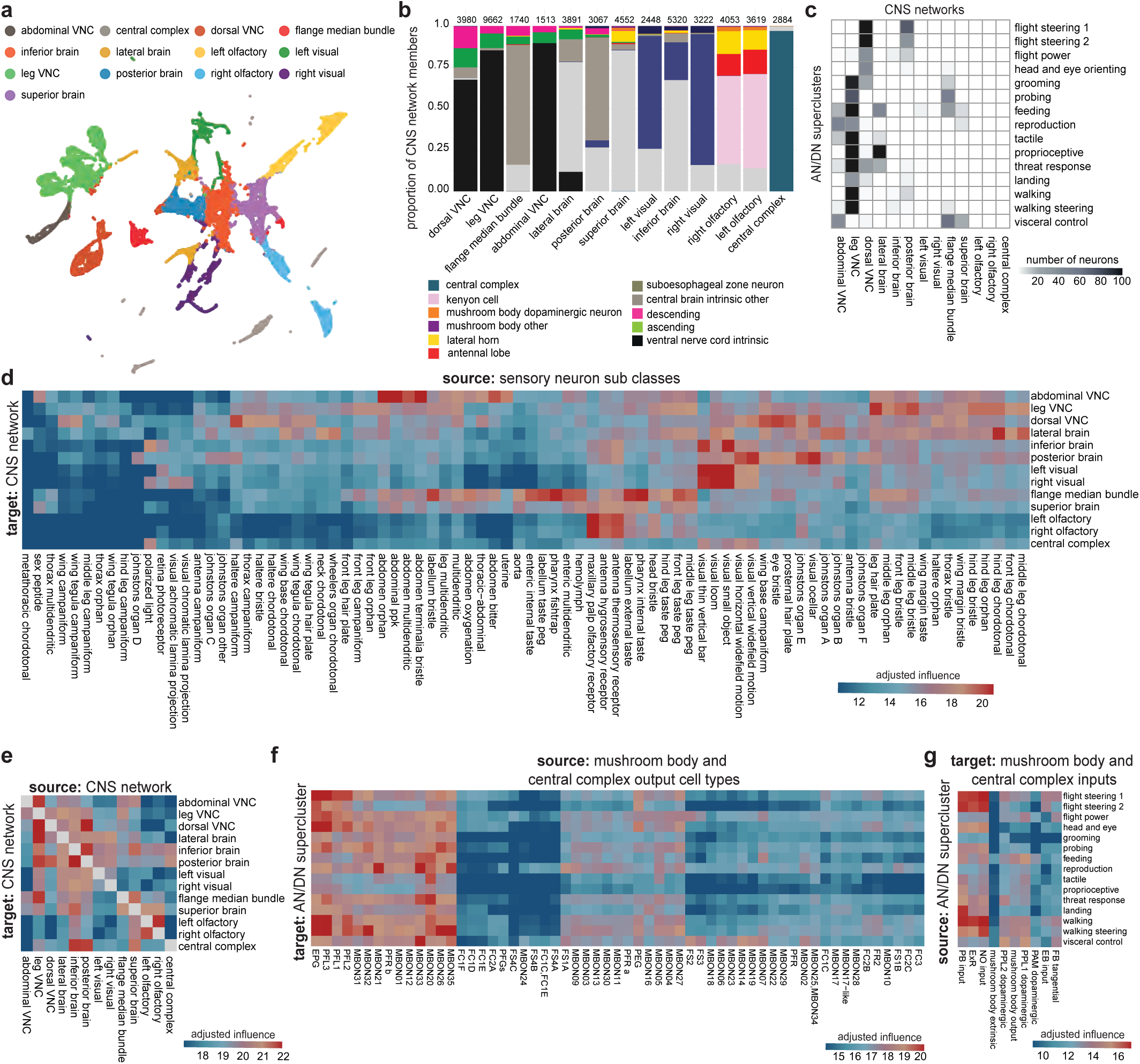
CNS networks’ cluster influence from sensors and to effectors. a. A UMAP embedding based on direct connectivity between BANC neurons (specifically, the eigenvectors of the graph Laplacian, derived from the undirected connectivity matrix), where each point is a neuron. This analysis uses all BANC neurons that meet four criteria: they are marked as proofread, they are intrinsic neurons of the CNS (not sensory or effector neurons), they have >100 incoming and outgoing connections, and no part of the cell is in the optic lobe (as the optic lobes are still undergoing proofreading). In total, 38394 neurons were used for this analysis, corresponding to 92% of cell-typed central brain and/or VNC intrinsic neurons. b. Proportion of each CNS network belonging to select super classes / cell classes. Numbers above each bar indicate the total number of neurons in each CNS network. c. Number of ANs and DNs in each CNS network. ANs and DNs are grouped by supercluster (Fig. 3d). d. Adjusted influence of sensors onto CNS networks. Visual projection neuron cell types are included, although they are not sensory neurons. e. Adjusted influence of each CNS network into each other CNS network. f. Adjusted influence of mushroom body output neurons and central complex output neurons onto AN/DN superclusters. g. Adjusted influence of AN/DN superclusters onto input neurons of the mushroom body and central complex.

## Supplementary Data

### Supplementary Data 1: Metadata categories and terms

Table of categories of annotations applied to BANC neurons and the list of terms used in each category. For region, side, flow, super_class, cell_class, cell_sub_class, cell_type, and hemilineage, only one term applies per neuron. For the other categories, neurons can be labeled with more than one term. Note that we do not include in this table all the possible options for cell_type, other_names, fafb_783_cell_type, manc_121_cell_type, fanc_1116_cell_type, hemibrain_121_cell_type, fafb_783_match_id, manc_121_match_id, fanc_1116_match_id, and hemibrain_121_match_id because there are too many possible options, but we include the categories and their descriptions to document them. For metadata annotations for each BANC neuron, see the data sources described in Data Availability.

- flow - from the perspective of the whole CNS, whether the neuron is afferent, efferent or intrinsic
- super_class - coarse division, hierarchical below flowcell_class - hierarchical below super_class
- cell_sub_class - hierarchical below cell_class
- cell_type - the name of the matched neuron from FAFB if it is a brain neuron or a DN or the name of the matched neuron from MANC if it is a VNC neuron or an AN. There are a few exceptions where those names did not define single cell types and were further split. This is hierarchical below cell_sub_class
- region - region of the CNS; all neurons that have arbors in the optic lobe are considered optic_lobe and all neurons that fully transit the neck connective between the brain and VNC are considered neck_connective
- side - from the fly’s perspective, the side on which the cell body is located or for afferent neurons, the side of the entry nerve.
- cell_function - term briefly describing the function of the neuron, applied largely to afferent and efferent neurons
- cell_function_detailed - more detailed information for the function of the neuron than cell_function, also applied largely to afferent and efferent neurons
- peripheral_target_type - the sensor or effector structure/organ targeted by an afferent/efferent neuron.
- body_part_sensory - the part of the body innervated by an afferent neuron
- body_part_effector - the part of the body targeted by an effector neuron. If known, this is the site of action when it is different from the body part innervated (e.g. wing power motor neurons innervate muscles located in the thorax but move the wing)
- nerve - peripheral nerve (if applicable)
- hemilineage - developmental lineage (NA for many neurons)
- sexually_dimorphic - isomorphic, dimorphic, or female-specific for whether the neuron is sexually dimorphic/sex-specific
- neurotransmitter_verified/neuropeptide_verified - neurotransmitter/neuropeptide of neuron, as reported in the literature
- neurotransmitter_predicted - CNN-predicted primary neurotransmitter
- other_names - names given to the neuron that are not the cell_type name
- fafb_783_cell_type/manc_121_cell_type/fanc_1116_cell_type/hemibrain_121_cell_type - cell type of neuron from FAFB v783/MANC v1.2.1/FANC v1116/Hemibrain v1.2.1 that matches the BANC neuron
- fafb_783_match_id/manc_121_match_id/fanc_1116_match_id/hemibrain_121_match_id - segment ID of neuron from FAFB v783/MANC v1.2.1/FANC v1116/Hemibrain v1.2.1 that matches the BANC neuron

### Supplementary Data 2: Updated annotations for FAFB Brain Neurons

Contains metadata for brain neurons from the FAFB-FlyWire dataset that are integrated into BANC analyses. This enables comparison and integration between the BANC neck connective data and the comprehensive adult brain connectome. Cell type names are unchanged.

- root_783 - FlyWire neuron ID (root_id in FAFB dataset release 783)
- nerve - peripheral nerve (if applicable)
- hemilineage - developmental lineage (NA for many neurons)
- region - region of the CNS; all neurons that have arbors in the optic lobe are considered optic_lobe and all neurons that fully transit the neck connective between the brain and VNC are considered neck_connective
- flow - from the perspective of the whole CNS, whether the neuron is afferent, efferent or intrinsic
- super_class - coarse division, hierarchical below flow
- cell_class - hierarchical below super_class
- cell_sub_class - hierarchical below cell_class
- cell_type - Individual cell type name (e.g., ORN_DM6, ORN_VA1v). Not modified from original project
- neurotransmitter_predicted - CNN-predicted primary neurotransmitter^51^
- neurotransmitter_verified - neurotransmitter, as reported in the literature

### Supplementary Data 3: Updated annotations for MANC VNC Neurons

Contains metadata for ventral nerve cord neurons from the MANC dataset that are integrated into BANC analyses. This enables comparison and integration between the BANC neck connective data and the comprehensive adult VNC connectomes. Cell type names unchanged.

- bodyid - MANC neuron ID from v1.2.1
- nerve - Peripheral nerve association (if applicable)
- hemilineage - Developmental lineage (NA for many neurons)
- region - region of the CNS; all neurons that have arbors in the optic lobe are considered optic_lobe and all neurons that fully transit the neck connective between the brain and VNC are considered neck_connective
- flow - from the perspective of the whole CNS, whether the neuron is afferent, efferent or intrinsic
- super_class - coarse division, hierarchical below flow
- cell_class - hierarchical below super_class
- cell_sub_class - hierarchical below cell_class
- cell_type - Individual cell type name (e.g., SNpp50, IN19A001). Not modified from original project
- neurotransmitter_predicted - CNN-predicted primary neurotransmitter
- neurotransmitter_verified - neurotransmitter, as reported in the literature

### Supplementary Data 4: ANs and DNs with UMAP coordinates and cluster assignments

Contains the ANs and DNs, along with their functional clustering based on connectivity patterns (**Fig. 3d**, **Extended Data Fig. 6**)

- root_id - BANC neuron identifier when used in analysis
- root_626 - BANC release v626 specific identifier
- supervoxel_id - supervoxel identifier for position
- position - 3D coordinates in BANC space (x, y, z in BANC raw voxel space)
- UMAP1, UMAP2 - 2D embedding coordinates from connectivity-based UMAP analysis
- side - from the fly’s perspective, the side on which the cell body is located
- region - region of the CNS (primarily neck_connective)
- nerve - peripheral nerve (if applicable)
- super_class - ascending, descending. Note, we only included flow == ‘intrinsic’ neurons
- hemilineage - developmental lineage
- cell_function - functional role description from our literature review
- cluster - cluster assignment from **Extended Data Fig. 6**. The number defines the cluster identity. Note that ANs have AN_ appended in front of the number and DNs have DN_ appended, but cells with the same number belong to the same cluster, regardless of the prefix
- super_cluster - AN/DN superclusters, the named cluster amalgamations used in this paper’s figures
- cell_type - BANC-specific cell type name, for DNs this preferentially comes from FAFB, for ANs from MANC
- fafb_cell_type - corresponding cell type in FAFB dataset
- manc_cell_type - corresponding cell type in MANC dataset

### Supplementary Data 5: effector neurons with UMAP coordinates and functional cluster assignments

Contains all efferent neurons, clustered by their functional properties and target effector systems (**Extended Data Fig. 4f**). These neurons control movement, secretion and other output functions.

- root_id - BANC neuron identifier when used in analysis
- root_626 - BANC release v626 specific identifier
- supervoxel_id - supervoxel identifier for position
- position - 3D coordinates in BANC space (x, y, z in BANC raw voxel space)
- UMAP1, UMAP2 - 2D embedding coordinates from connectivity-based UMAP analysis
- side - from the fly’s perspective, the side on which the cell body is located
- region - region of the CNS
- nerve - peripheral nerve
- super_class - efferent type (motor, visceral_circulatory)
- cell_class - hierarchical below super_class, incorporates innervated body part
- cell_sub_class - hierarchical below cell_class, incorporates innervated body part and some function
- hemilineage - developmental lineage
- body_part_effector - the part of the body targeted by an effector neuron. If known, this is the site of action when it is different from the body part innervated (e.g. wing power motor neurons innervate muscles located in the thorax but move the wing)
- cell_function - functional role (e.g. leg_motor, antenna_motor, neck_motor).
- cluster - cluster assignment from **Extended Data Fig. 4f**, as the cluster number with EFF_ appended (e.g., EFF_01)
- super_cluster - effector neuron groups as the named clusters used in **Extended Data Fig. 4f** and subsequent figures.
- cell_type - BANC-specifìc cell type name
- fafb_cell_type - corresponding cell type in FAFB dataset
- manc_cell_type - corresponding cell type in MANC dataset

### Supplementary Data 6: CNS network analysis with spectral clustering and UMAP embedding

Contains neurons from spectral clustering analysis of the CNS connectivity (**Fig. 6a**), revealing network-level organisation beyond individual cell types. This analysis identifies functional networks that span multiple brain regions.

- root_id - BANC neuron identifier when used in analysis
- root_626 - BANC release v626 specific identifier
- supervoxel_id - supervoxel identifier for position
- position - 3D coordinates in BANC space (x, y, z in BANC raw voxel space)
- UMAP1, UMAP2 - 2D embedding coordinates from connectivity-based UMAP analysis
- side - from the fly’s perspective, the side on which the cell body is located
- region - region of the CNS
- nerve - peripheral nerve (if applicable)
- super_class - high-level functional category (various types including visual_projection, central_brain_intrinsic)
- hemilineage - developmental lineage
- cell_function - functional description (if known)
- cluster - effector neuron clusters (from **Extended Data Fig. 4f**), which have the EFF_ prefix, and AN/DN clusters (from **Extended Data Fig. 6**), which have the AN_ or DN_ prefix (if applicable)
- super_cluster - name of effector neuron group or AN/DN supercluster (if applicable)
- cns_network - CNS networks as determined by spectral clustering, 13 cluster cut
- cell_type - BANC-specific cell type name
- fafb_cell_type - corresponding cell type in FAFB dataset
- manc_cell_type - corresponding cell type in MANC dataset

### Supplementary Data 7: Literature review on cell function for ascending, descending and visual projection neurons

- cell_type - cell type names in the BANC connectome
- other_names - other names used for this cell type in the literature
- super_class - high-level functional category, here only ascending, descending and visual projection
- cell_function - simple descriptive label for the ‘function’ of the cell type
- citations - short hand citations for the work that helped determine cell_function

### Supplementary Data 8: Bounding boxes for known dataset artefacts that negatively impact neuronal reconstruction

Bounding boxes delineating regions with known data-quality issues in the BANC dataset. Coordinates are in BANC raw-voxel space (1 voxel = 4 × 4 × 45 nm).

- issue - short informal label that names the issue type
- min_x, min_y, min_z - lower corner of the bounding box (voxels)
- max_x, max_y, max_z - upper corner of the bounding box (voxels)

